# Surface-Based Connectivity Integration

**DOI:** 10.1101/2020.07.01.183038

**Authors:** Martin Cole, Kyle Murray, Etienne St-Onge, Benjamin Risk, Jianhui Zhong, Giovanni Schifitto, Maxime Descoteaux, Zhengwu Zhang

## Abstract

There has been increasing interest in jointly studying structural connectivity (SC) and functional connectivity (FC) derived from diffusion and functional MRI. However, several fundamental problems are still not well considered when conducting such connectome integration analyses, e.g., “Which structure (e.g., gray matter, white matter, white surface or pial surface) should be used for defining SC and FC and exploring their relationships”, “Which brain parcellation should be used”, and “How do the SC and FC correlate with each other and how do such correlations vary in different locations of the brain?”. In this work, we develop a new framework called *surface-based connectivity integration* (SBCI) to facilitate the integrative analysis of SC and FC with a re-thinking of these problems. We propose to use the white surface (the interface of white matter and gray matter) to build both SC and FC since diffusion signals are in the white matter while functional signals are more present in the gray matter. SBCI also represents both SC and FC in a continuous manner at very high spatial resolution on the white surface, avoiding the need of pre-specified atlases which may bias the comparison of SC and FC. Using data from the Human Connectome Project, we show that SBCI can create reproducible, high quality SC and FC, in addition to three novel imaging biomarkers reflective of the similarity between SC and FC throughout the brain, called global, local, and discrete *SC-FC coupling*. Further, we demonstrate the usefulness of these biomarkers in finding group effects due to biological sex throughout the brain.

## 1. Introduction

Brain connectivity is an intriguing and quickly expanding research field [Eyewire and Blake, 1997; Park and Friston, 2013; Smith et al., 2015; Glasser et al., 2016b; Shi and Toga, 2017]. With recent advances in magnetic resonance imaging (MRI) techniques, we are able to noninvasively probe the human brain at higher resolutions than ever before [Glasser et al., 2016a] and construct different types of connectomes. Among them, two imaging modalities, diffusion MRI (dMRI) and functional MRI (fMRI), and their corresponding brain connectivities are particularly prominent. While dMRI measures the restriction of isotropic diffusive water movement [Basser et al., 1994] and can be used to infer structural connectivity (SC) [Sporns, 2013b], fMRI measures the blood oxygen level dependent (BOLD) signal, deemed as a proxy for neurovascular coupling due to large-scale neural activation [Ogawa et al., 1990] and can be used to infer functional connectivity (FC) [Sporns, 2013a].

Both SC and FC play important roles in understanding the human brain [Sporns, 2013a; Zimmermann et al., 2018]. As a result of the anatomy of WM, dMRI can be used to estimate the restricted diffusive patterns of water throughout the brain by applying magnetic gradients of multiple strengths in many directions [Li et al., 2016]. Diffusion signals are then reconstructed per voxel and represented as fiber orientation distribution functions (fODFs), which can then be fitted to tractography algorithms to estimate the locations of WM connections [Basser et al., 1994; Descoteaux et al., 2009; Bastiani et al., 2012; Tournier et al., 2012; Girard et al., 2014]. SC can be constructed from the tractography results [Zhang et al., 2018]. On the other hand, FC is obtained from calculating the correlation of BOLD signals between different brain regions. Differences in magnetic susceptibilities between oxygenated and deoxygenated blood give rise to the BOLD signal after neural stimulation has occurred in a region [Ogawa et al., 1990]. BOLD imaging produces a time series of these susceptibility changes and is thought to represent the overall blood oxygenation exchange in each voxel as a function of time.

Although most existing connectivity studies explore SC or FC independently, there is increasing interest in exploring SC and FC together, referred to as SC and FC integration in this paper. Much of this previous integration work roughly falls into three broad classes: *prediction, modeling*, and *fusion*. As one of the earliest papers studying the relationships between SC and FC, Honey et al. [2009] demonstrated that brain regions that are directly structurally connected have higher FC than those regions that are not directly connected, leading to a series of studies that attempted to *predict* an individual’s FC directly from SC [Honey et al., 2010; Goñi et al., 2014; Messé et al., 2015; Deligianni et al., 2013; Chamberland et al., 2017; Higgins et al., 2018; Li et al., 2019]. *Modeling* FC with neural spiking models by incorporating known SC as prior information has also drawn some attention recently. By assuming that SC and FC are highly correlated, it may be possible to derive FC patterns directly from neural spiking equations and SC to simulate hemodynamic response functions (HRFs) without collecting any functional data [Iyer et al., 2013; Nakagawa et al., 2013; Bassett et al., 2018; Wang et al., 2019]. Finally, connectivity *fusion* uses measurements from both dMRI and fMRI to derive new information about the brain [Zhu et al., 2014; Bassett et al., 2018]. For example, Fan et al. [2016] built a novel brain atlas using information from SC and FC, incorporating both structural and functional information across the brain.

These previous studies of SC and FC integration used different image resolutions, tractography algorithms, and brain parcellations to reconstruct the connectomes. Some found that the strength of FC is related to the anatomical pathways (SC) [Honey et al., 2009; Van Den Heuvel et al., 2009; Bowman et al., 2012; Hermundstad et al., 2013; Goñi et al., 2014], while others concluded that the correlation between SC and FC is poor [Ghumman et al., 2016; Buckner et al., 2013; Chamberland et al., 2017]. These heterogeneous findings lead to several fundamental questions we must consider when studying SC and FC jointly:

1. Which structure (e.g., gray matter, white matter, the white or pial surface) is the best place in the brain to explore the relationships between SC and FC?
2. Which parcellation is most suitable for constructing and comparing SC and FC?
3. What is the correlation or coupling strength between SC and FC throughout the brain and how does it vary across different populations?

Most existing studies use gray matter (GM) regions of interest (ROIs) as network nodes in deriving SC and FC. However, due to the limitations in dMRI acquisition [Reveley et al., 2015] and streamline reconstruction [Thomas et al., 2014; Girard et al., 2014; Maier-Hein et al., 2017], using GM ROIs to build SC can result in biased SC estimation. For example, streamlines can stop prematurely in the WM or near the GM-WM interface, superficial WM tracts can impede the construction of longer streamlines [Reveley et al., 2015], and the precision of streamline reconstruction is reduced due to partial volume effects (PVEs) in lower spatial resolution acquisitions [Tournier et al., 2011]. Similarly, using GM ROIs as nodes in FC often requires volumetric smoothing and PVE correction, which reduces the spatial localization of BOLD signals and thus provides inaccurate FC estimation [Coalson et al., 2018].

Traditional construction of SC and FC often requires predefined parcellations/atlases for two primary reasons: 1) dimensionality reduction and 2) more straightforward interpretations of localized physical processes [Glasser et al., 2016a]. However, choosing an optimal brain parcellation for SC and FC integration is challenging due to the complicated nature of the brain. Parcellations derived from one modality [Gordon et al., 2014; Thomas Yeo et al., 2011] may not be suitable for studying others. As such, we currently do not know the most suitable parcellation to study SC and FC jointly.

Finally, the correlation or coupling strength between SC and FC has the potential to unlock some key insights on how brain structure collaborates with function since it links both dMRI and fMRI, where each modality provides us with crucial information to better understand the underlying brain connectivity. For example, it is natural to expect that such correlation can be spatially different across brain regions and populations [Nakagawa et al., 2013]. However, few studies have evaluated the relationship between SC and FC at high-resolution, nor how this relationship may differ between different groups of subjects (e.g., males vs. females).

With a comprehensive re-thinking of these problems, we propose a new pipeline, named Surface-Based Connectivity Integration (SBCI), to facilitate the integrative analysis of SC and FC. In SBCI, we propose to use the white surface (the interface of white matter and gray matter) to build both SC and FC and represent them in a continuous fashion without the need of any pre-specified brain parcellation. SBCI also gives three novel SC-FC coupling strength measures, two of which have not been studied before to the best of our knowledge. Figure 1 shows a systematic overview of the SBCI framework. Compared with existing work, SBCI has the following unique features:

**Figure 1:**
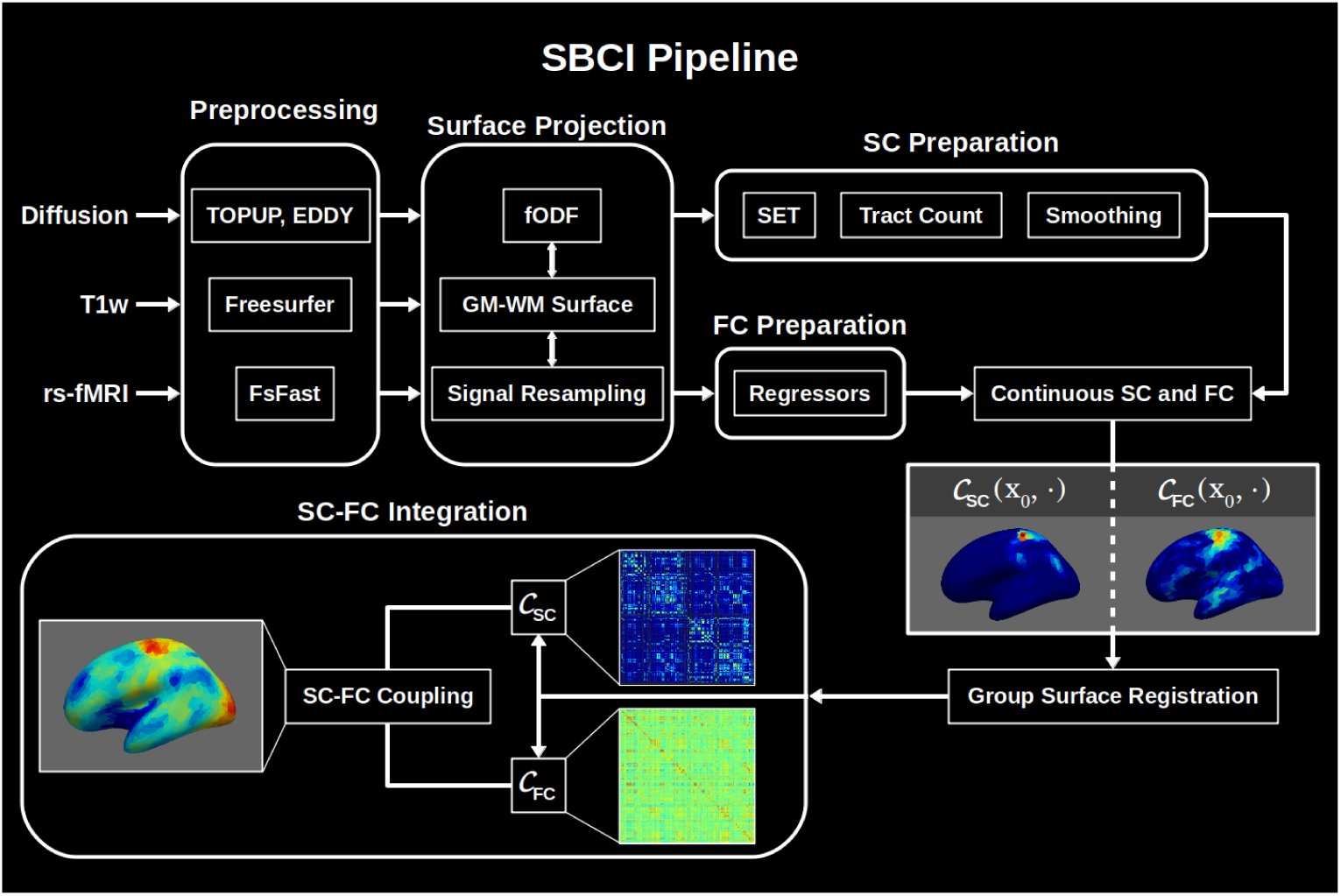
Flowchart of the SBCI pipeline. SBCI: surface-based connectivity integration; SC: structural connectivity; FC: functional connectivity; 𝒞_*SC*_ : continuous SC; 𝒞_*F C*_ : continuous FC; SET: surface-enhanced tractography; FsFast: Freesurfer’s Functional Analysis Stream; GM: gray matter; WM: white matter; fODF: fiber orientation distribution function.

- **SBCI maps SC and FC to the white surface.** The white surface is the interface between cortical GM and WM. Since diffusion signals are mostly in the WM regions and functional signals are more present in the GM regions, we posit that the white surface is the best place to study the integration of SC and FC. The HCP team promotes using brain surfaces to study FC due to better functional signal localization and more straightforward surface registrations between subjects [Glasser et al., 2016a; Coalson et al., 2018]. However, surface-based SC studies are still rare in the literature. Extending tractography reliably to the white surface comes with many challenges [Reveley et al., 2015]. A novel algorithm called surface-enhanced tractography (SET) [St-Onge et al., 2018] overcomes many of these challenges by incorporating prior knowledge from the geometry of the white surface. Instead of using the unreliable dMRI signal near the white surface, SET uses a surface flow technique to model the superficial WM structure. In SET, all reconstructed streamlines intersect with the white surface.
- **SBCI treats SC and FC as continuous functions.** For any two points on the white surface, SBCI defines connectivity strengths for SC and FC. We smooth the sparse SC to result in a continuous measure [Moyer et al., 2017] that facilitates integration with FC. Continuous treatment of SC and FC has two clear advantages in the study of SC-FC integration: (1) connectomes can be represented at very high spatial resolution compared to current atlas-based methods and (2) there is no need to rely on predefined templates or parcellations to construct connectivity matrices, allowing for a more flexible and robust treatment of connectivity analysis [Messé, 2019].
- **SBCI produces continuous and discrete SC-FC coupling features on the white surface.** We define SC-FC coupling (SFC) as the similarity between continuous SC and FC at any point on the white surface. SBCI gives three measures of different SFC features: continuous global coupling between SC and FC without a predefined parcellation, continuous local coupling within ROIs defined by a given parcellation, and discrete coupling with a given parcellation. Details of these SFC definitions are presented in Section 2.5.

In the following sections, we describe the details of the SBCI pipeline, experiments used to validate it, and an application of the SFC features as imaging biomarkers to distinguish biological sex.

## 2. Methods

### 2.1. Datasets

The HCP intiative is the current gold standard for human brain connectivity mapping. For this study, in order to develop and validate the SBCI pipeline, we use the HCP Young Adult (HCPYA) and HCP Test-Retest (HCPTR) data. Data are downloaded from the ConnectomeDB website.

The data used in this paper include preprocessed T1-weighted (T1w) and dMRI images and unprocessed resting-state fMRI images from the HCPYA and HCPTR datasets. Full imaging acquisition information and minimal fMRI and dMRI image preprocessing steps are documented in Glasser et al. [2013]. Briefly, all imaging was conducted on the 3T Siemens Connectom scanner (Erlangen, Germany). High-resolution T1w anatomical images were acquired with the 3D MPRAGE (magnetization prepared rapid gradient echo) sequence with a slice acceleration factor of 2 using 0.7 mm isotropic resolution. Diffusion imaging was performed using a 2D spin-echo EPI (echo planar imaging) sequence with approximately 90 diffusion directions at three non-zero b-values (1,000, 2,000, and 3,000 s/mm^2^ each) and 6 *b*0 reference scans at 1.25 mm isotropic resolution. A full diffusion MRI run includes 6 runs of about 9 mins 50 seconds each, representing 3 gradient tables, with each table acquired once with right-to-left (RL) and left-to-right (LR) phase encoding polarities, respectively. Resting-state functional imaging was performed using a 2D gradient-echo EPI sequence with repetition time 720 ms, echo spacing 33.1 ms, and 2 mm isotropic resolution. Parallel imaging was enabled using a multi-band acceleration factor of 8. Resting state fMRI scans were acquired in 4 runs of approximately 15 minutes each, with eyes open in a dark room. Runs alternated encoding polarities, resulting in two RL and two LR scans.

Our HCPTR data includes 38 subjects with complete MR imaging data collected at Washington University in St. Louis as a follow-up to an initial HCPYA scan, resulting in two full sets of imaging data (test and retest) for each subject. Our HCPYA data includes 89 random subjects (46 females) from the S500 data release in the 26 − 30 year old group. We also tested SBCI in a few subjects collected in a Siemens MAGNETOM PrismaFit (Erlangen, Germany) scanner at the University of Rochester to verify that SBCI can give valid results in relatively low-resolution MRI data (details about the data and SBCI results are presented in Supplementary Material Section 1).

### 2.2. Image Preparation and Preprocessing

Diffusion image preprocessing steps, including brain extraction, susceptibility induced distortion correction, motion correction, and eddy-current distortion correction, are performed using tools in FMRIB’s software language (FSL) [Woolrich et al., 2009; Smith et al., 2004; Sotiropoulos et al., 2013]. For both HCP datasets described above, we download the diffusion data after eddy-current correction. We begin with the T1w anatomical images after gradient correction and the raw rs-fMRI. Additional diffusion processing is performed using a combination of tools from Mrtrix3 [https://www.mrtrix.org/], advanced normalization tools (ANTs) [Avants et al., 2011], and the Sherbrooke Connectivity Imaging Lab toolbox in Python (Scilpy), including resampling to 1 mm isotropic resolution and building the fiber orientation distribution functions (fODFs) to prepare for tractography.

The anatomical T1w processing includes registration to the high-resolution diffusion images via ANTs and surface reconstruction using the recon all tool available in Freesurfer [http://freesurfer.net/].

Resting-state fMRI images are processed using Freesurfer’s functional analysis stream (FsFast) and includes motion correction, brain masking, sampling to the surface (left and right), and surface smoothing with a Gaussian kernel with FWHM *σ* in Freesurfer. Common nuisance variables are calculated in Freesurfer, including the WM signal, CSF signal, six motion parameters (three translational and three rotational), and the global signal [Murphy and Fox, 2017], where we include the top 5 principal components (PCs) for WM and CSF signals. More details about this regression can be found in Murphy et al. [2013]; Fox et al. [2009] and Murphy and Fox [2017]. Additional processing details are available in Supplemental Material Section 2.

### 2.3. Continuous Functional Connectivity on the White Surface

During the FsFast pipeline, we map the volumetric BOLD signals to the subject’s 32k white surface, resulting in a BOLD time series at each vertex on the surface meshes. The FC between any pair of vertices is calculated using a partial correlation between the two BOLD time series [Kirch, 2008], controlling for confounding signals.

Let 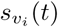 be the residual BOLD time series after regressing out nuisance variables at the *i*-th vertex *v*_*i*_. The FC between any vertex pair (*v*_*i*_, *v*_*j*_) is calculated as:

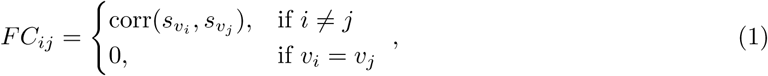

where we do not consider self-interactions.

Although we can get a very high-resolution FC with dense white surface meshes, the FC obtained in (1) is still considered to be discrete. In our SBCI pipeline, we represent FC in a continuous fashion. Hence, here we introduce the continuous FC concept with some mathematical formalism. We use the convention that a vertex is a location on the mesh grid, and a point is any location on the surface. For any pair of points on the white surface, we want to evaluate their FC and therefore assume there is a BOLD time series at every point on the white surface.

More specifically, let Ω be the union of two disjoint white surfaces of the brain; the continuous FC can be represented as a symmetric function on the space Ω × Ω. For any pair of points (*x, y*) ∈ Ω × Ω, we define the continuous FC as 𝒞_*FC*_ (*x, y*) = corr(*s*_*x*_, *s*_*y*_), where *s*_*x*_ and *s*_*y*_ are the BOLD time series at points *x* and *y*. We define the set of all possible FCs as ℱ_*FC*_ = {𝒞_*FC*_ : Ω × Ω → [−1, 1] : 𝒞_*FC*_ (*x, y*) = 0 if *x* = *y*; and 𝒞_*FC*_ (*x, y*) = 𝒞_*FC*_ (*y, x*)}.

Note that there are two major problems in this continuous FC framework. First, we only observe BOLD signals at discrete vertices, not every point on Ω. Second, the white surfaces between subjects are different, making any group-wise analyses (analyses across multiple subjects) difficult. To solve these problems we inflate each white surface into two 2-spheres, and with some abuse of notation also denote Ω as the union of two 2-spheres 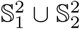. The geometry of a 2-sphere will more easily facilitate signal processing and inter-subject alignment [Dale et al., 1999; Fischl et al., 1999a].

To obtain a BOLD signal for any point *x* ∈ Ω, we apply the following signal interpolation method on the 2-sphere. Without loss of generality, assume 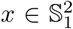 (one 2-sphere in Ω), let *B*(*x*; *σ*) represent a neighborhood near *x* such that all vertices in *B*(*x*; *σ*) have geodesic distances to *x* less than *σ*. We have the BOLD time course *s*_*x*_ at *x* calculated as *s*_*x*_ = ∑_*v*′∈*B*(*x*;*σ*)_ *w*_*σ*_(*x, v*′;) ∗ *s*_*v*_, where *w*_*σ*_(*x, v*′) can be a truncated Gaussian or biweight (quartic) kernel with bandwidth *σ* defined with a geodesic distance on the 2-sphere, e.g., *w*_*σ*_(*x, v*′) = (15*/*16*σ*){1 − (*d*(*x, v*′)*/σ*)^2^}^2^𝕀_*d*(*x,v*′)<*σ*_ [Risk and Zhu, 2019].

When multiple subjects are involved in the analysis, registration between subjects is necessary, e.g., finding correspondence among Ω_1_, …, Ω_*n*_ for *n* subjects. One of the advantages of inflating the original complex white surfaces to two 2-spheres is that registration of signals on a 2-sphere is much easier than on an irregular manifold space [Coalson et al., 2018; Glasser et al., 2016a; Robinson et al., 2014; Kurtek et al., 2010]. We defer the registration between subjects to Section 2.5.2.

### 2.4. Continuous Structural Connectivity on the White Surface

#### 2.4.1. Extending Streamlines to the White Surface

SC aims to measure the extent to which brain regions are connected by WM fiber bundles, where traditionally regions are obtained based on a parcellated atlas [Zhang et al., 2018, 2019a]. In the context of image analysis, we are limited to inferring SC from the streamlines connecting any pair of brain regions constructed by tractography algorithms [Thomas et al., 2014; Girard et al., 2014; Maier-Hein et al., 2017] due to MRI acquisition limitations, such as low spatial resolution and signal-to-noise ratio. While different measures of SC have been proposed to characterize the structural connection between two regions [Zhang et al., 2018], for simplicity we choose to study SC using streamline count. Once all streamlines are constructed throughout the brain, we naively define the discrete SC as the number of streamlines between any two regions.

We use the white surface to construct SC, which requires that all reconstructed streamlines intersect with the surface. Unfortunately, most tractography algorithms cannot project streamlines to the surface due to two primary challenges. First, diffusion signals are unreliable in the GM and near the GM-WM interface regions due to the anatomy of neurons, leading to an overestimation of streamlines in the WM that do not intersect the white surfaces [Girard et al., 2014; Reveley et al., 2015]. Second, there will be gyral biases in SC as shown in St-Onge et al. [2018] due to a few factors, including low spatial resolutions during acquisition and biased tractography seeding choice.

A recent development called surface-enhanced tractography (SET) [St-Onge et al., 2018] overcomes these challenges, allowing streamlines to be reliably extended to the white surface. In SBCI, we apply SET to build streamlines. The white surface is used to initiate the flow with a parameter *t* controlling for the amount of flow into the WM, resulting in a surface beneath the white surface. Starting from the surface at *t* > 0, we initialize streamlines using *N*_*s*_ seed points on the surfaces and propagate them using the particle filtering technique (PFT) [Girard et al., 2014]. Figure 2 illustrates some results from the SET pipeline, where (a) shows the initial white matter segments constructed by SET and (b) shows the final tractography result. Experiments have shown that SET decreases SC gyral biases and better approximates the underlying anatomy by using a more stringent assumption in regions where dMRI signals are less informative [St-Onge et al., 2018].

**Figure 2:**
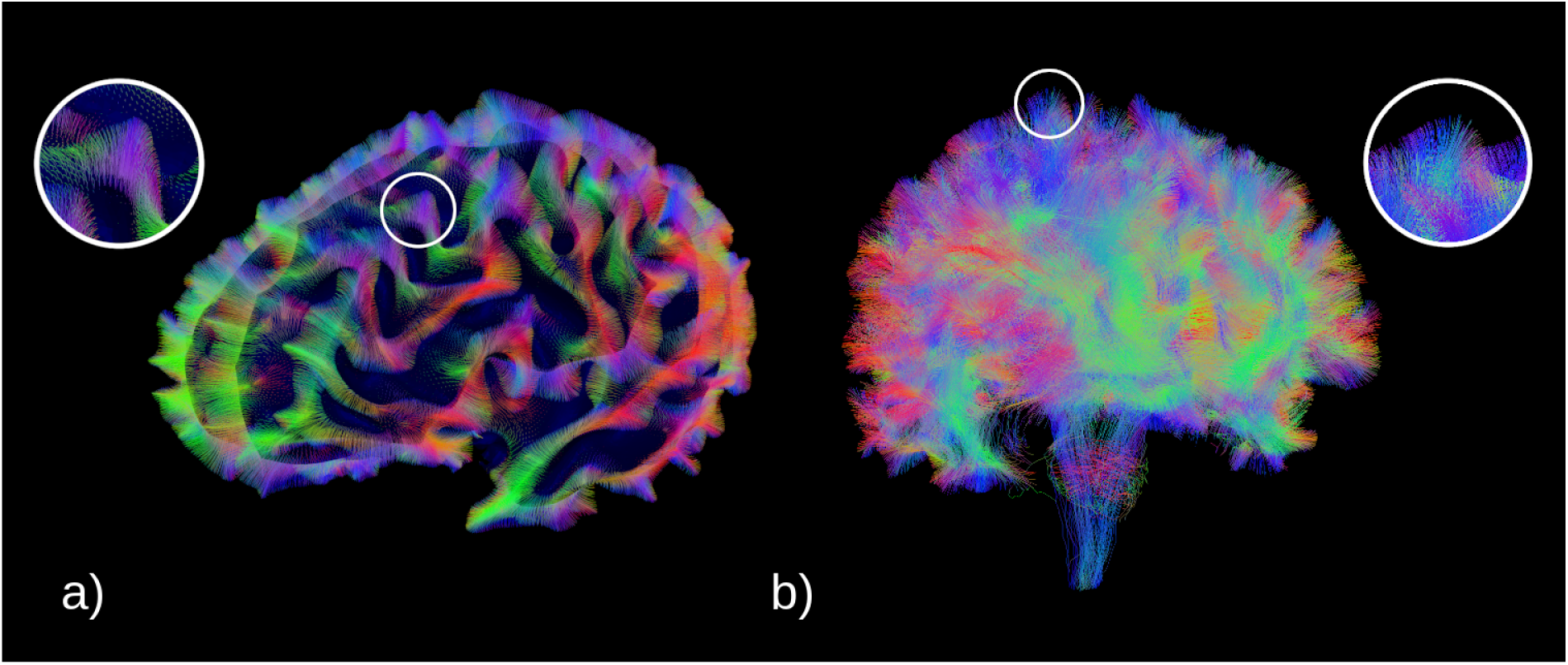
Tractography results from SET. (a) shows the surface flow to the white surface, and (b) shows the final tractography with the reconstructed fanning structure near the white surface.

#### 2.4.2. Smoothing for Continuous Structural Connectivity

In this paper we deviate from the traditional parcellation-based approach to SC and instead define a continuous framework (similar to FC as described in Section 2.3). Maintaining both SC and FC in a comparable continuous framework allows for a more robust treatment of the integration of the two modalities. For any two distinct points *x* and *y* on the brain surfaces, we define SC as 𝒞_*SC*_ (*x, y*) ∈ ℝ_+_ ∪ {0}, where SC is the probability density function representing the likelihood that *x* and *y* are structurally connected by WM fibers. 𝒞_*SC*_ is a symmetric function defined on Ω × Ω and the set of all SC functions can be denoted as ℱ_*SC*_ = {𝒞_*SC*_ : Ω × Ω ↦ ℝ_+_ ∪ {0} : 𝒞_*SC*_ (*x, y*) = 0 if *x* = *y*; 𝒞_*SC*_ (*x, y*) = 𝒞_*SC*_ (*y, x*) and ∬_Ω×Ω_ 𝒞_*SC*_ (*x, y*)*dxdy* = 1}. Note that the continuous SC has been proposed before in different formats and applications, e.g., in Moyer et al. [2017] and Gutman et al. [2014].

The limited number of streamlines (typically a few million) constructed in Section 2.4.1 with SET gives us a discrete and sparse sampling of 𝒞_*SC*_ (refer to Figure 4 panel (a)). In contrast, FC has non-zero values for nearly every connection. In order to more fairly compare and integrate structural and functional connectivities, we propose to estimate a smooth and dense SC using kernel density estimation (KDE) on Ω × Ω. Assume that we have a symmetric kernel *k*_*h*_ defined as a mapping from Ω × Ω to ℝ_+_ ∪ {0} with a bandwidth parameter *h*. The smoothed SC under a standard KDE procedure is given by:

**Figure 3:**
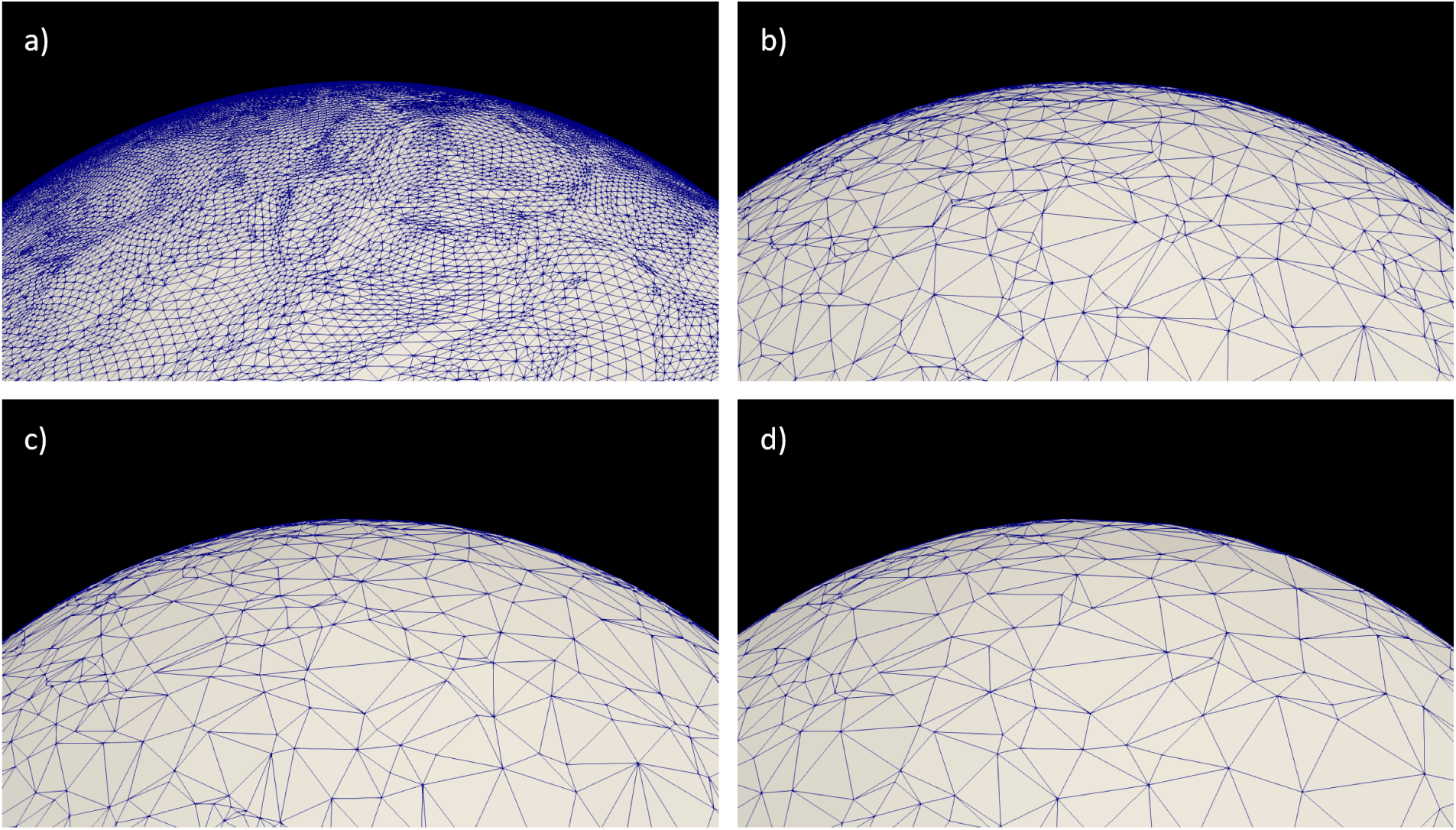
Example grids of Ω in SBCI. From (a) to (d), we have different sparsity levels of vertices on a 2-sphere: (a) 163,842 vertices; (b) 8453 vertices; (c) 5157 vertices; and (d) 1834 vertices. The down-sampling is conducted by minimizing the Hausdorff distance between the full and lower resolution meshes.

**Figure 4:**
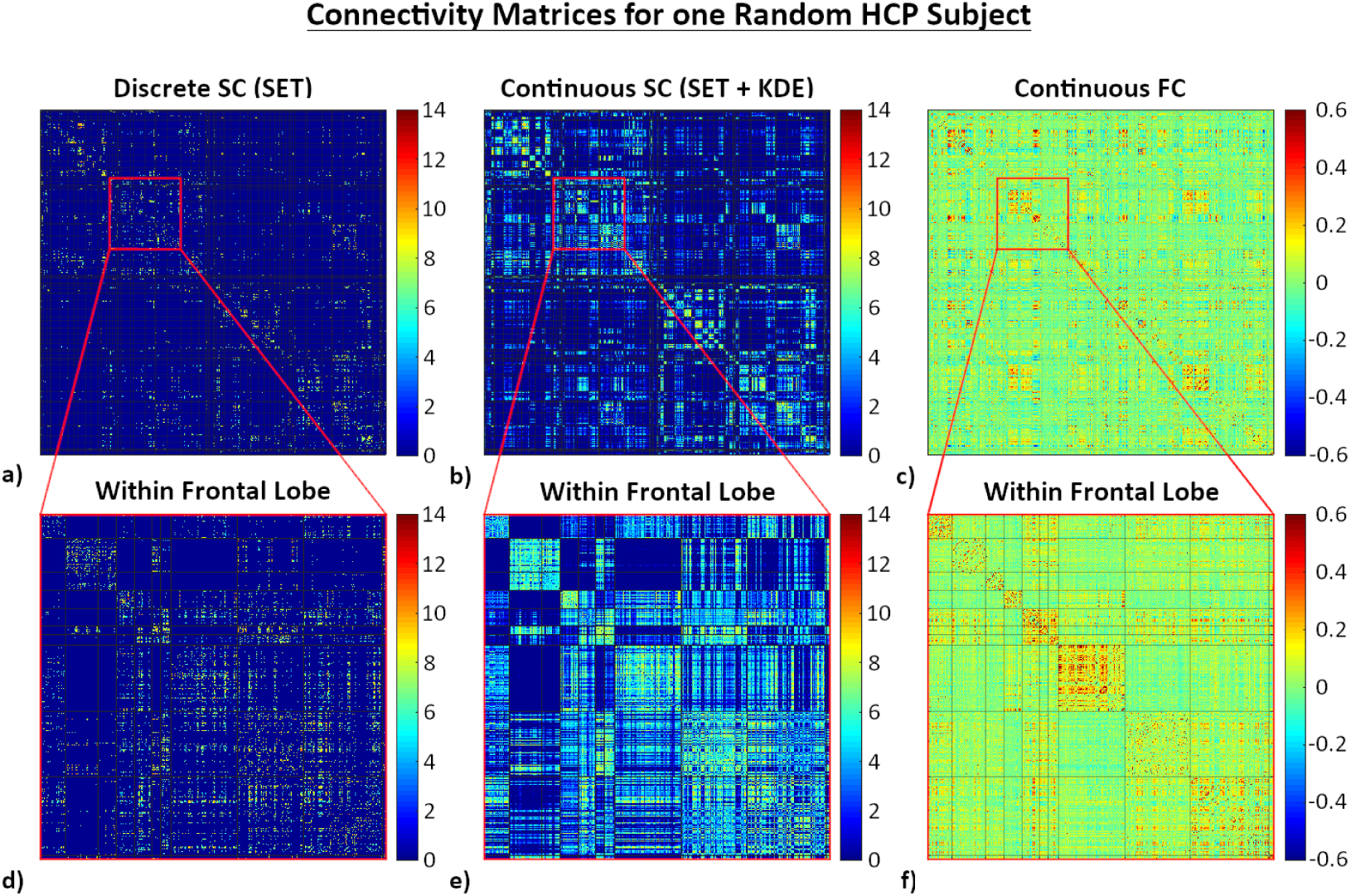
Outputs from SBCI. (a) Discrete SC before smoothing; (b) Continuously smoothed SC using *h* = 0.005; (c) Continuous FC, and (d-f) zoomed in on the left frontal lobe for connectomes in (a-c). The black horizontal and vertical lines in (e-f) designate different ROIs in the Desikan-Killiany atlas for comparison with typical atlas-based connectivity matrices.

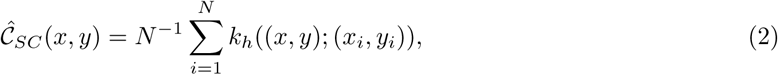

where *N* is the total number of observed streamlines, and (*x*_*i*_, *y*_*i*_) represents the endpoints of the *i*-th observed streamline.

To use the KDE method, we must define an appropriate kernel function *k*_*h*_ on the domain Ω × Ω. Similar to Moyer et al. [2017] and Risk and Zhu [2019], we begin by defining a symmetric heat kernel [Hartman and Watson, 1974] on a 2-sphere as

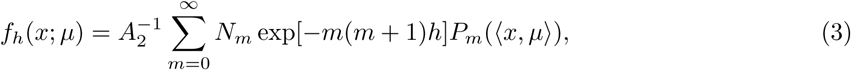

where *µ* ∈ 𝕊^2^ and *h* ∈ ℝ_+_ represent the mean and bandwidth, respectively, *A*_2_ = 4*π* (the area of 𝕊^2^), *m*(*m* + 1) are the eigenvalues of the Laplacian on 𝕊^2^ for *m* = 0, 1, …, ∞, *P*_*m*_ is the Legendre polynomial of order *m* for ℝ^3^, *N*_*m*_ equals 2*m* + 1, the number of linearly independent homogeneous spherical harmonics of degree *m* in ℝ^3^, and ⟨, ⟩ indicates the inner product of two elements on 𝕊^2^.

Since 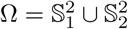, we extend *f*_*h*_ to Ω trivially by letting *f*_*h*_(*x*; *µ*) = 0 if *x* and *µ* are not on the same sphere. We then define a kernel function on the domain Ω × Ω using a product of two functions *f*_*h*_ on Ω. Given a mean (*µ*_*x*_, *µ*_*y*_) the kernel is defined as *k*_*h*_((*x, y*); (*µ*_*x*_, *µ*_*y*_)) = *f*_*h*_(*x, µ*_*x*_)*f*_*h*_(*y, µ*_*y*_).

The bandwidth of the kernel function is a key parameter under KDE. Although many bandwidth selection criterion have been proposed [Bowman, 1984; Turlach, 1993; Jones et al., 1996b,a; Botev et al., 2010; Moyer et al., 2017; Zhang et al., 2019b; Risk and Zhu, 2019], there is no consensus on the best general approach. In this paper, we select *h* based on the reproducibility of 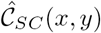 calculated on the HCPTR dataset.

### 2.5. SC-FC Coupling

With continuous SC and FC defined on the domain Ω × Ω, we are ready to study the integration of structural and functional connectivities on the white surface for an individual or a group of subjects. We first describe three different kinds of SC-FC coupling (SFC) at the subject level. Then we describe the alignment procedure needed for studying SFC at the group level.

#### 2.5.1. SC-FC Coupling at the Subject Level

##### Continuous Global SC-FC Coupling

We first evaluate the consistency between SC and FC on the surface without any predefined parcellation of the brain. Let 𝒞_*SC*_ (*x, y*) and 𝒞_*FC*_ (*x, y*) denote the continuous SC and FC for a particular subject. At any point *x*_0_ in Ω, we define the global SC-FC coupling using a normalized inner product of two functions *f*_1_(*y*) = 𝒞_*SC*_ (*x*_0_, *y*) and *f*_2_(*y*) = 𝒞_*FC*_ (*x*_0_, *y*):

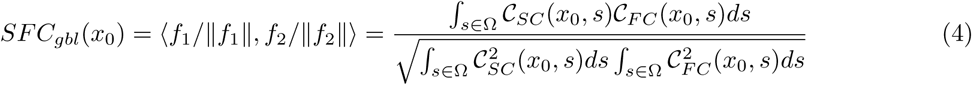

The inner product ⟨,⟩ is defined for two functions on Ω, which is analogous to calculating the correlation between two rows of a functional connectivity and structural connectivity matrix in a traditional setting.

The *SFC*_*gbl*_(*x*_0_) returns a scalar at *x*_0_, and therefore, *SFC*_*gbl*_ is a continuous function on Ω, measuring the similarity/consistency between SC and FC at different locations on the white surface. In practice, we evaluate the *SFC*_*gbl*_ on a discrete grid of Ω. The mesh surfaces from Freesurfer provide a natural choice for such a grid. However, the dense vertices on these surfaces cause computational challenges. In our implementation of SBCI, we down-sample each white surface mesh (left and right) from over 120,000 vertices to around 2100 using the Visualization Toolkit (VTK) in Python [Schroeder et al., 2006, 1992]. To maintain as much topological information as possible, we sample vertices from the white surface such that the induced Hausdorff distance [Aspert et al., 2002] between the full and down-sampled meshes is minimized. We then generate the down-sampled white, inflated, and spherical surface meshes using Delaunay triangulation [Barber et al., 1996] and the coordinates corresponding to the sampled vertices on the full meshes. Figure 3 shows a surface mesh before (panel (a)) and after down-sampling at three different sparsity levels of vertices (panels (b), (c) and (d)). With the downsampled mesh in (d), in the experiment section, our final continuous SC and FC are represented with matrices of 3668 × 3668 dimensions before masking out the corpus callosum region.

##### Continuous Local SC-FC Coupling

When a parcellation of the brain surface is available, we can evaluate the coupling strength of our continuous SC and FC within ROIs. For an ROI *E* and a point *x*_0_ ∈ *E*, our continuous local SFC (*SFC*_*loc*_) is defined as:

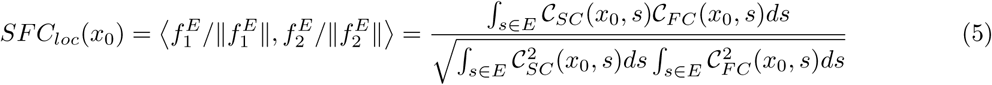

To the best of our knowledge, within-region connectivity correlation has not been explored before in the literature.

##### Discrete Parcellation-based SC, FC and SC-FC Coupling

At last, following existing literature [Baum et al., 2020; Cocchi et al., 2014; Wang et al., 2018; Jiang et al., 2019; Zhang et al., 2011], we define discrete SFC for a given parcellation. We first convert our continuous SC and FC to finite adjacency matrices based on the given parcellation. For any two ROIs *E*_1_ and *E*_2_, we define the SC between the two regions as:

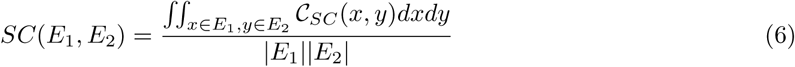

where |*E*_*i*_| represents the area of region *E*_*i*_ for *i* = 1, 2. The defined SC strength represents the connectivity density in a unit area square.

The traditional way to calculate FC is to first obtain a mean BOLD signal for each region and then calculate the Pearson correlation coefficient. To be more consistent with the SC calculation, we instead consider an average of the correlations. We calculate the FC in the following way: first apply the Fisher z-transformation to the correlation, calculate the average, and then apply the inverse Fisher z-transformation:

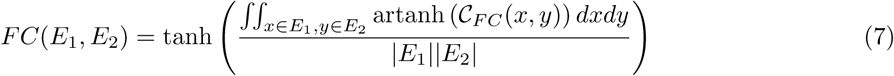

where tanh(·) and artanh(·) represent the hyperbolic tangent function and its inverse, respectively. Finally, we define the discrete SC-FC coupling (*SFC*_*dct*_) as:

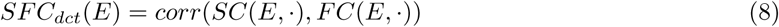

where *corr*(·, ·) represents the Pearson correlation, and *SC*(*E*, ·) and *FC*(*E*, ·) represent the structural and functional connections between ROI *E* and all other ROIs, respectively.

#### 2.5.2. SC-FC Coupling at the Group Level

At the individual level, there is no need to perform alignment between subjects. However, different brains have different surfaces with varying shapes and sizes; thus, registration between subjects is necessary when comparing the continuous SFC features between subjects. One advantage of registration on the 2-sphere compared to 3D volumetric registration is that the 2D sphere has one less dimension than the 3D volume of the brain, which can improve registration accuracy [Glasser et al., 2016a]. To register brain surfaces between subjects, we find correspondence between Ω_1_, …, Ω_*n*_ from *n* subjects according to an objective function [Fischl et al., 1999b; Robinson et al., 2014]. Note that here we consider each of the two spheres in Ω independently. Without loss of generality, we consider alignment between two subjects and let *g*_1_, *g*_2_ : 𝕊^2^ ↦ ℝ^*k*^ be feature functions of the two subjects being aligned. Registering *g*_1_ to *g*_2_ is the process of finding a mapping function (also called a warping function) *γ* from 𝕊^2^ to 𝕊^2^ such that some objective function ℒ(*g*_1_(*γ*(·)), *g*_2_(·)) is minimized, where *γ* ∈ Γ and Γ is the set of all diffeomorphisms of 𝕊^2^ to itself [Kurtek et al., 2010].

The Freesufer software [Fischl et al., 1999a,b] performs alignment by using the feature function *g* generated by the folding pattern of the cortical surface and matching this folding pattern between subjects. More advanced algorithms have also been proposed by using more features derived from multiple MRI modalities [Glasser et al., 2016a; Robinson et al., 2014]. In this paper, we use the alignment algorithm and registration results from Freesurfer. More specifically, for each subject we have two mapping functions (*γ*_1_, *γ*_2_), one for each hemisphere, to align the subject to a template (e.g., fsavarage space). Let 𝒞_*SC*_ (*x, y*) and 𝒞_*FC*_ (*x, y*) be the subject’s continuous SC and FC. We have the aligned SC and FC represented as 𝒞_*SC*_ ((*γ*_1_, *γ*_2_)(*x*), (*γ*_1_, *γ*_2_)(*y*)) and 𝒞_*FC*_ ((*γ*_1_, *γ*_2_)(*x*), (*γ*_1_, *γ*_2_)(*y*)), where (*γ*_1_, *γ*_2_)(*x*) indicates that if *x* is on 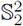, we have (*γ*_1_, *γ*_2_)(*x*) = *γ*_1_(*x*) and if *x* is on 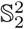, we have (*γ*_1_, *γ*_2_)(*x*) = *γ*_2_(*x*). After alignment, we can calculate SC-FC coupling features on a discrete grid of Ω, which can then be compared across subjects.

### 2.6. Evaluation and Analysis

To construct and validate the SBCI pipeline, we perform the following analyses. Note that in our experiments, two parcellation approaches are used: the Desikan-Killiany [Desikan et al., 2006] atlas and Atlas-Free (the continuous SC, FC, and SFC). Experiments in the parameter selection include the Destrieux [Destrieux et al., 2010] atlas as well.

#### 2.6.1. SBCI Parameter Selection

A few parameters in SBCI are critical to connectome mapping: the surface flow size *t* in SET, the number of tractography seeds *N*_*s*_, and the SC smoothing bandwidth *h*. Using the HCPTR dataset, we optimize these parameters based on the reproducibility of our final SC. The reproducibility is measured using the distance-based intraclass correlation coefficient (dICC) defined in Zhang et al. [2018], which is a generalization of the intraclass correlation coefficient (ICC) [Shrout and Fleiss, 1979], bounded by 1. Higher dICC values indicate better reproducibility.

The surface smoothing kernel FWHM *σ* is another parameter in SBCI that can be tuned for the functional data. We select *σ* = 5 *mm* to remain consistent with typical fMRI preprocessing procedures.

#### 2.6.2. SBCI Connectome Reproducibility

To validate the SBCI pipeline, we perform qualitative and quantitative exploratory analyses to assess the reliability of our pipeline and compare them to previous studies. These analyses fall into three categories: SC reproducibility, FC reproducibility, and the relationship between SC and FC. Comparisons are made using the Desikan-Killiany atlas and Atlas-Free approach. More specifically, we perform visual inspections of SC and FC at different spatial resolutions (different grids on Ω, refer to Figure 3) and calculate the ICC and dICC with the HCPTR dataset. We also compare the distributions of FC strength with and without direct SC connections. We calculate the ICC values for all connections that have nonzero SC values between nodes in all subjects in the HCPTR dataset (to avoid the zero-inflation problem [Shrout and Fleiss, 1979]). The dICC is calculated under the Frobenius norm using the entire connectivity matrix.

#### 2.6.3. SC-FC Coupling Reproducibility

Upon confirming the validity of our SBCI pipeline to produce consistent and reproducible SC and FC at both standard atlas and high resolutions, we seek to examine the reproducibility of the SFC features.

After registering each subject’s images to the standard fsaverage space, we obtain aligned SC and FC on a common white surface. Using the SFC definitions presented in Section 2.5 and the Desikan-Killiany atlas, we quantify the reproducibility of each feature using both ICC and dICC measures. The four features considered in this paper are: (1) global SC-FC coupling (*SFC*_*gbl*_ - Eq. (4)); (2) local SC-FC coupling based on the the Desikan-Killiany atlas (*SFC*_*locdk*_ - Eq. (5)); (3) local SC-FC coupling using major brain lobes (*SFC*_*loclb*_ - Eq. (5)) and (4) discrete SC-FC coupling based on the Desikan-Killiany atlas (*SFC*_*dct*_ - Eq. (8)). Note that for *SFC*_*loclb*_ the lobar parcellation was created by merging ROIs within the Desikan-Killiany atlas to achieve 12 large areas (right and left frontal, parietal, temporal and occipital lobes, and the right and left insula and cingulate).

To more directly show the intra- and inter-subject differences on the SFC feature, we conduct the following analysis. In the HCPTR data, we have one dMRI and four rs-fMRI runs in each of two scanning sessions, allowing us to compare SFC features measured from the same subject at different time points and from different subjects. At each *x* ∈ Ω, we perform paired t-tests to determine: (1) SFC differences across time points (test and retest) within one subject and (2) SFC differences between different subjects. We expect more significant differences (smaller p-values) to be present between subjects than across time points within single subjects.

#### 2.6.4. SC-FC Coupling as Biomarkers

In order to assess the usefulness of the SFC features as imaging biomarkers, we use the HCPYA dataset, described in section 2.1, to find group differences due to sex. This study is motivated by past findings that male and female brains have anatomical, functional, and biochemical differences [Zaidi, 2010; Weis et al., 2020]. We also conduct a classification experiment to test and compare the discriminative ability of different connectivity features (continuous SC, FC, and these SFC features) in distinguishing males from females. Principle component analysis (PCA) is applied to reduce each feature’s dimensionality before different classifiers are used and evaluated with five fold cross-validation, where we used the logistic regression classifier (LRC), support vector classifier (SVC), and random forest classifier (RFC). Receiver operator characteristic (ROC) areas under the curve (AUCs) are used to evaluate classification performance.

## 3. Results

### 3.1. Parameter Selection

Here we present results for the selection of three important parameters in SCBI: the flow parameter *t*, the number of seeds *N*_*s*_, and the KDE bandwidth *h*. Using the HCPTR data, we calculated the dICC using different values of *t* (*t* ∈ {75, 100}) and *N*_*s*_ (*N*_*s*_ ∈ {1.5 ∗ 10^6^, 3 ∗ 10^6^, 4.5 ∗ 10^6^, 6 ∗ 10^6^, 7.5 ∗ 10^6^}) for three different parcellation approaches: Desikan-Killiany, Destrieux, and Atlas-Free. These results are shown in Supplementary Material Tables 1 and 2. We find that the flow parameter *t* = 75 produces higher dICC values, and thus more reproducible SC, for both the Destriuex and Atlas-Free approaches regardless of the number of seeds used. Additionally, for all approaches, dICC increases as a function of the number of seeds used, but this increase does not account for the increased computational power needed when using more seeds. Therefore, we select *t* = 75 and *N*_*s*_ = 3 ∗ 10^6^ in all subsequent experiments. Using a 2.70GHz Intel CPU it takes around 4.5 hours to to generate half a million streamlines per 1.5 million seeds used.

**Table 1:** Reproducibility measure dICC vs. bandwidth for the continuous SC. The table shows results when using three atlases (after removing outliers) with different KDE bandwidth parameters (*h*).

| Atlas | $h = 0$ | $h = 0.002$ | $h = 0.005$ | $h = 0.010$ | $h = 0.020$ |
| --- | --- | --- | --- | --- | --- |
| Desikan | 0.7706 | 0.8054 | **0.8062** | 0.8046 | 0.7999 |
| Destrieux | 0.7798 | 0.7791 | 0.7910 | 0.7983 | **0.7991** |
| Atlas-Free | 0.7501 | 0.7873 | **0.7954** | 0.7945 | 0.7877 |

**Table 2:** Sex prediction mean_±*stdev*_ AUC scores for different classification models with 5-fold stratified cross-validation. LRC=Logistic Regression, SVC=Support Vector Classifier, RFC=Random Forest Classifier. K corresponds to the number of principal component scores used in these models.

| Model | $N_{PCs}$ | Continuous SC | Continuous FC | $SFC_{gbl}$ | $SFC_{locdk}$ | $SFC_{loclb}$ | $SFC_{dct}$ |
| --- | --- | --- | --- | --- | --- | --- | --- |
| LRC | K=5 | $0.75 \pm 0.14$ | $0.66 \pm 0.11$ | $0.71 \pm 0.08$ | $0.74 \pm 0.11$ | $0.71 \pm 0.07$ | $0.71 \pm 0.16$ |
| | K=10 | $0.86 \pm 0.01$ | $0.79 \pm 0.09$ | $0.81 \pm 0.05$ | $0.86 \pm 0.12$ | $0.85 \pm 0.09$ | $0.72 \pm 0.18$ |
| | K=20 | $0.81 \pm 0.07$ | $0.80 \pm 0.07$ | $0.79 \pm 0.18$ | $0.81 \pm 0.12$ | $0.77 \pm 0.13$ | $0.68 \pm 0.14$ |
| SVC | K=5 | $0.76 \pm 0.13$ | $0.54 \pm 0.16$ | $0.70 \pm 0.03$ | $0.71 \pm 0.11$ | $0.71 \pm 0.05$ | $0.57 \pm 0.16$ |
| | K=10 | $0.85 \pm 0.02$ | $0.78 \pm 0.07$ | $0.83 \pm 0.05$ | $0.84 \pm 0.10$ | $0.87 \pm 0.14$ | $0.75 \pm 0.18$ |
| | K=20 | $0.80 \pm 0.06$ | $0.79 \pm 0.06$ | $0.83 \pm 0.10$ | $0.79 \pm 0.11$ | $0.78 \pm 0.12$ | $0.70 \pm 0.15$ |
| RFC | K=5 | $0.70 \pm 0.14$ | $0.60 \pm 0.11$ | $0.61 \pm 0.07$ | $0.60 \pm 0.15$ | $0.67 \pm 0.13$ | $0.68 \pm 0.12$ |
| | K=10 | $0.82 \pm 0.08$ | $0.65 \pm 0.19$ | $0.69 \pm 0.08$ | $0.76 \pm 0.15$ | $0.79 \pm 0.10$ | $0.71 \pm 0.14$ |
| | K=20 | $0.86 \pm 0.06$ | $0.79 \pm 0.06$ | $0.72 \pm 0.11$ | $0.78 \pm 0.09$ | $0.69 \pm 0.08$ | $0.75 \pm 0.08$ |

The supplemental material includes additional parameter selection results before removing two outlier subjects from the HCPTR data (Supplementary Material Table 1). We determine these outliers by observing large differences in the test and retest surface meshes produced in the anatomical preprocessing steps. Removing the two outliers increases the dICC score for all approaches.

Next we explore an optimal choice for *h*. Table 1 shows dICC values for the three approaches with five different KDE bandwidth values, *h* ∈ {0, 0.002, 0.005, 0.01, 0.02} after removing two outliers. As our pipeline is designed to conduct connectivity analyses using an Atlas-Free approach, we select *h* = 0.005 because it maximizes the dICC value when using the Desikan-Killiany and Atlas-Free approaches. We see that smoothing is especially helpful for enhancing the reproducibility of high-resolution SC. In fact, after smoothing, the dICC for our Atlas-Free approach is nearly the same as the Desikan-Killiany and Destrieux parcellations, which is a substantial improvement over the original unsmoothed SC (*h* = 0). Using a 2.70GHz Intel CPU it takes approximately 8, 4, and 2 hours to smooth SC matrices obtained using grids Ω of 16,906, 10,314 and 3668 vertices, respectively, producing SC matrices requiring 1.1GB, 0.5GB and 0.1GB of storage. The full SBCI pipeline takes approximately 4-5 days (10 hours for fMRI processing, 1 day for T1w processing, 2-3 days for dMRI processing, and 12 hours for connectome integration) to run for a single subject.

### 3.2. Exploratory Connectome Analyses

With these selected parameters, we explore the brain connectomes (SC and FC) produced by SBCI. Figure 4 shows (a) discrete SC constructed at the same resolution as our continuous SC by SET, (b) continuous SC smoothed using *h* = 0.005, (c) continuous FC, and (d)-(f) are zoomed in on the left frontal lobe of the connectomes in (a)-(c). Note that we multiply a constant to the continuous SC in (b) for visualization purposes. Comparing (a) and (b), we can see that after KDE smoothing, we have a much denser SC, more similar to the FC, making the comparison between SC and FC more robust. Moreover, SC and FC matrices are traditionally constructed using discrete brain parcellations. By significantly increasing the spatial resolution of SC and FC, we can observe more detailed relationships between them, as shown in Figure 4 (e) and (f), which cannot be observed using atlas-based connectivity analyses. More results from the HCP and our local data are presented in Supplementary Material Figure 2 and Figure 4.

Next, we evaluate the relationship between SC and FC. In Figure 5, we show the histograms of FC strengths with and without direct SC connections. In our continuous SC framework, we define a direct SC connection as those node-pairs with SC values greater than 10^−7^. This means if ten million streamlines are built for a subject, the connections that have more than one expected streamline will be considered to have a direct SC connection. One advantage of our high-resolution SC and FC is that we can inspect such relationships in local brain regions as well as the whole brain [Honey et al., 2009]. From Figure 5, we see that in general, FC connection strengths are higher with direct SC connections compared to those without direct SC connections. The mean differences between the two types of FC are different for different brain regions, e.g., the left and right parietal lobes (left two plots in the first row) have larger differences than the left and right frontal lobes (left two plots in the second row). More interestingly, we see that FC strengths without direct SC connections have a mean strength close to zero, while FC strengths with direct SC connections have a mean strength greater than zero. These patterns are consistently observed in all subjects in our study.

**Figure 5:**
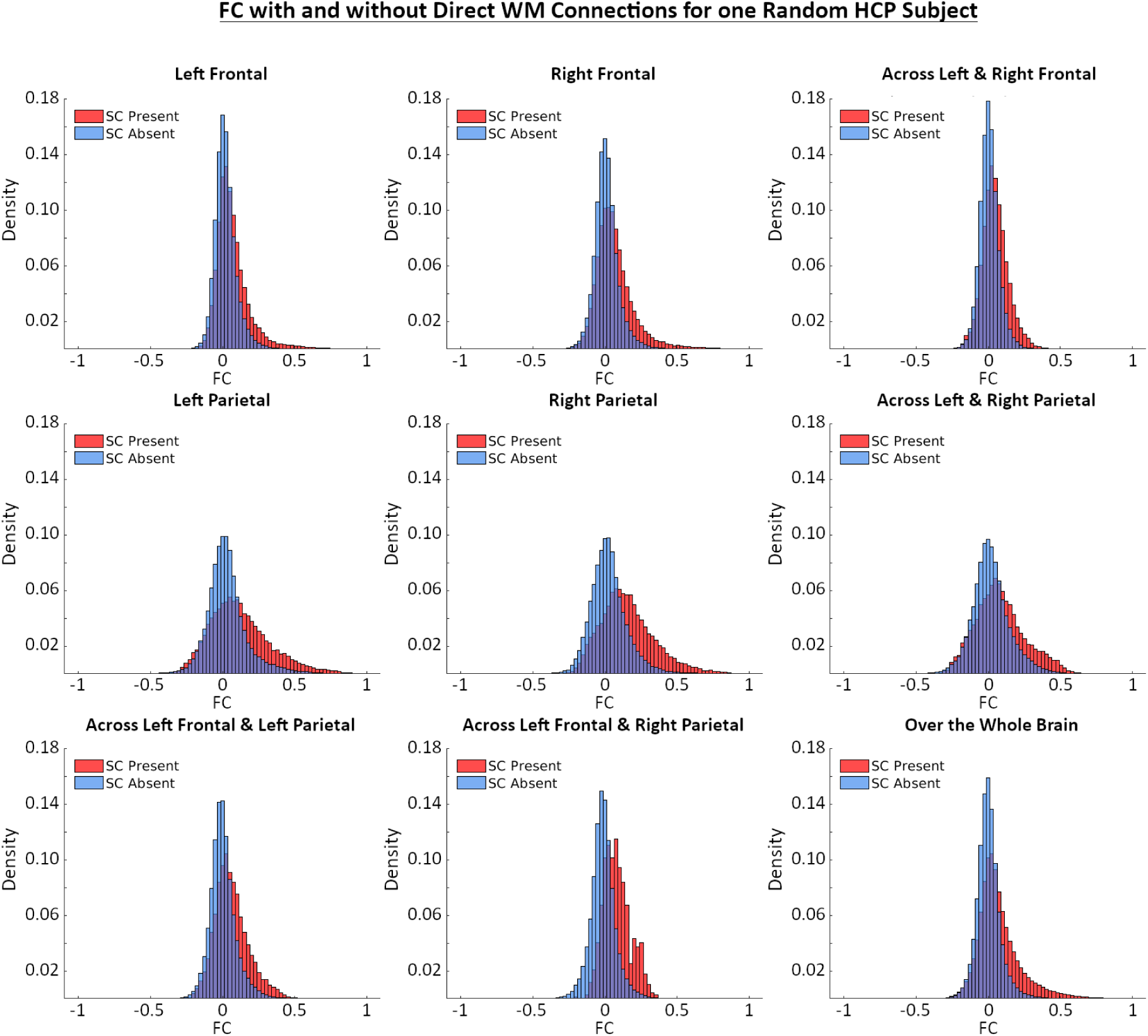
Histograms of resting state FC for nodes-pairs with and without direction WM SC connections. Within ROI histograms are calculated using all node-pairs within that region (defined by the Desikan-Killiany atlas), and across region histograms are calculated using node-pairs that are in different regions.

Finally, we quantitatively assess the reproducibility of SC and FC produced by SBCI. We compare the dICC and node-wise ICCs between two parcellation approaches: Desikan-Killiany and Atlas-Free. In each test and retest scanning session in the HCPTR dataset, we have one dMRI scan and four resting state fMRI runs, resulting in a total of 72 SC matrices and 288 FC matrices for the entire dataset. We place scans from the same subject next to each other, so that we can observe a block pattern along the diagonal of these distance matrices. Panels (a) and (b) in Figure 6 show the pairwise distance matrices for the structural and functional connectomes using the HCPTR dataset (after removing the same two outliers as the data producing Table 1) under the Desikan-Killiany and Atlas-Free approaches, respectively. Panels (c) and (d) show the connection-wise ICC distributions corresponding to the plots in panels (a) and (b). dICC values are displayed above each distance matrix. The mean and standard deviation of the ICC distributions are displayed above the accompanying histograms.

**Figure 6:**
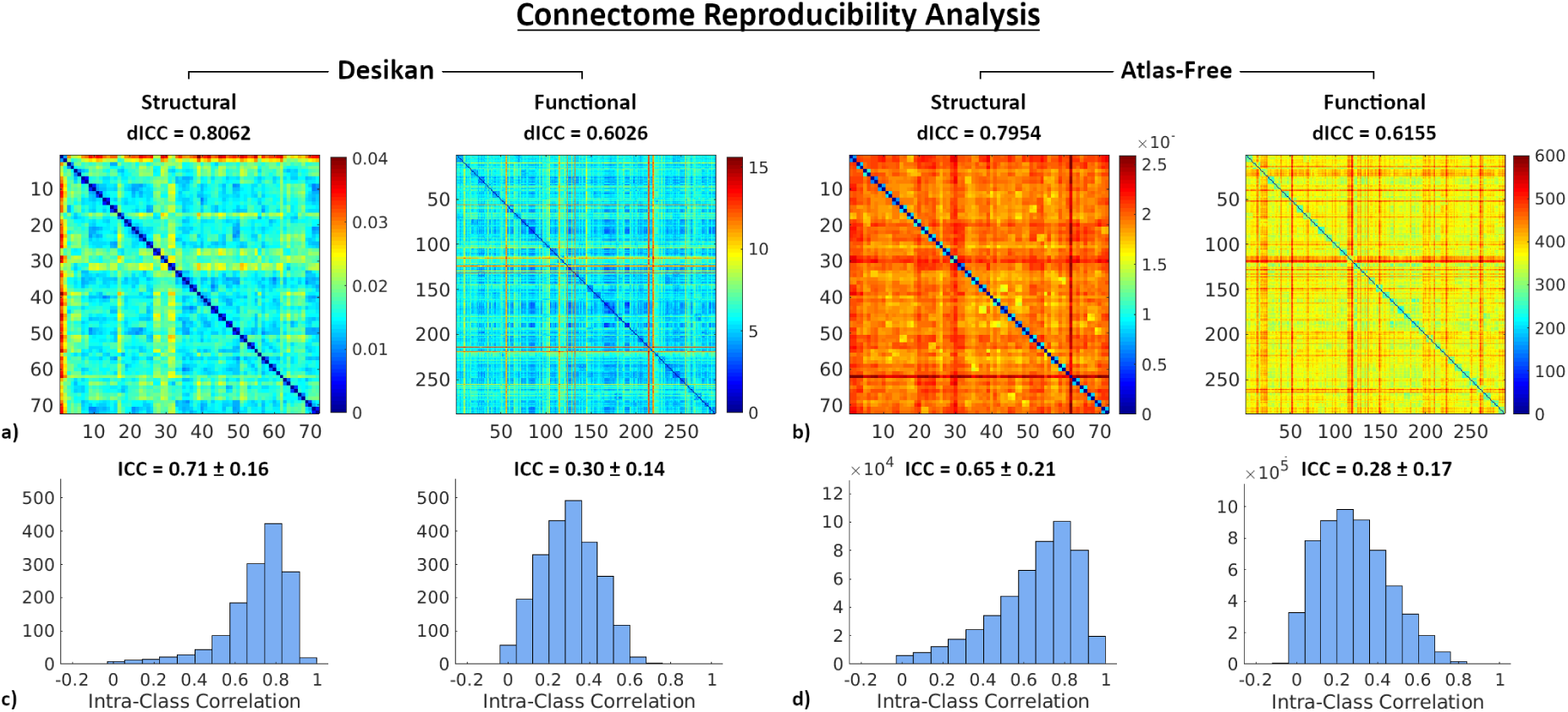
Reproducibility analysis of SC and FC from SBCI. The first row shows pairwise distance matrices of SCs and FCs generated under (a) the Desikan-Killiany atlas and (b) our Atlas-Free approach. We use the Frobenius norm to calculate the distance between connectivity matrices, and the different scales are due to the differences in the connectivity matrices generated by each method (the relative difference between the intra-subject distances and the inter-subject distances is more important than the magnitude of those distances). Note that we have 72 SCs and 288 FCs in the HCPTR dataset. dICC values are displayed above each distance matrix. Panels c) and d) show ICC histograms for every node-pair in the SCs and FCs corresponding panels a) and b), respectively. Mean and standard deviation are displayed above each histogram.

The dICC values of SCs derived from the Desikan-Killiany and Atlas-Free approaches are 0.8062 and 0.7954, respectively. The FC dICC values of the Desikan-Killiany and Atlas-Free approaches are 0.6026 and 0.6155, respectively. The maintenance of dICC values across both parcellation approaches for structural and functional data demonstrates that the within-subject variations of each connectome are less than between-subject variations. We also examine the reproducibility of individual connections by calculating node-wise ICC values. Structural ICC distributions have means (±standard deviations) 0.71±0.16 and 0.65±0.21 for the Desikan-Killiany and Atlas-Free approaches, respectively. Functional ICC distributions have mean (±standard deviations) 0.30±0.14 and 0.28±0.17 for the Desikan-Killiany and Atlas-Free parcellations.

In general, we observe mostly good (0.75 ≥ *ICC* > 0.6) to excellent (*ICC* > 0.75) SC reproducibility for both the Desikan-Killiany parcellation and our Atlas-Free framework. Also, our high-resolution Atlas-Free connectome reproducibility measured by the dICC is nearly on par with the Desikan-Killiany atlas-based connectome, indicating that SBCI can produce high-quality high-resolution SC. FC has poor (0.4 ≥ *ICC*) to fair (0.6 ≥ *ICC* > 0.4) reproducibility, with a majority of connections falling in the relatively poor reproducibility range. Despite relatively poor reproducibility, our FC reproducibility is consistent with those presented in the literature using the same dataset [Tomasi et al., 2017] and other processing pipelines [Noble et al., 2017]. Compared with previous reproducibility studies [Noble et al., 2019], SBCI maintains a typical FC reliability range even at very high spatial resolution.

### 3.3. SC-FC Coupling Reproducibility

We now evaluate the reproducibility of our SFC features. For each subject in the HCPTR dataset, we have 2 SC matrices and 4 FC matrices, and therefore obtain 8 (2 ∗ 4) SFC measures for each definition of SFC for each subject, resulting in 288 (8 ∗ 36) total features for all 36 subjects. Figure 7 shows the pairwise distance matrices, dICC, and ICC histograms for the four different SFC features. The dICC scores of the *SFC*_*gbl*_, *SFC*_*locdk*_, *SFC*_*loclb*_, and *SFC*_*dct*_ are 0.682, 0.728, 0.707, and 0.606, respectively. Most of the ICC values of the global and local SFC features fall in the fair to good reliability range. Comparing results in Figure 7 with the ones in Figure 6, we have a few interesting findings: (a) our SFC features have much better reproducibility values (as measured by both ICC and dICC) compared to the FC, especially the continuous SFC features; (b) the continuous local SFCs are more reproducible than the global SFC; and (c) the discrete SFC (*SFC*_*dct*_) is the least reproducible SFC feature. Plots of these SFC features for an individual can be found in Supplementary Material Figures 3, 5, and 6.

**Figure 7:**
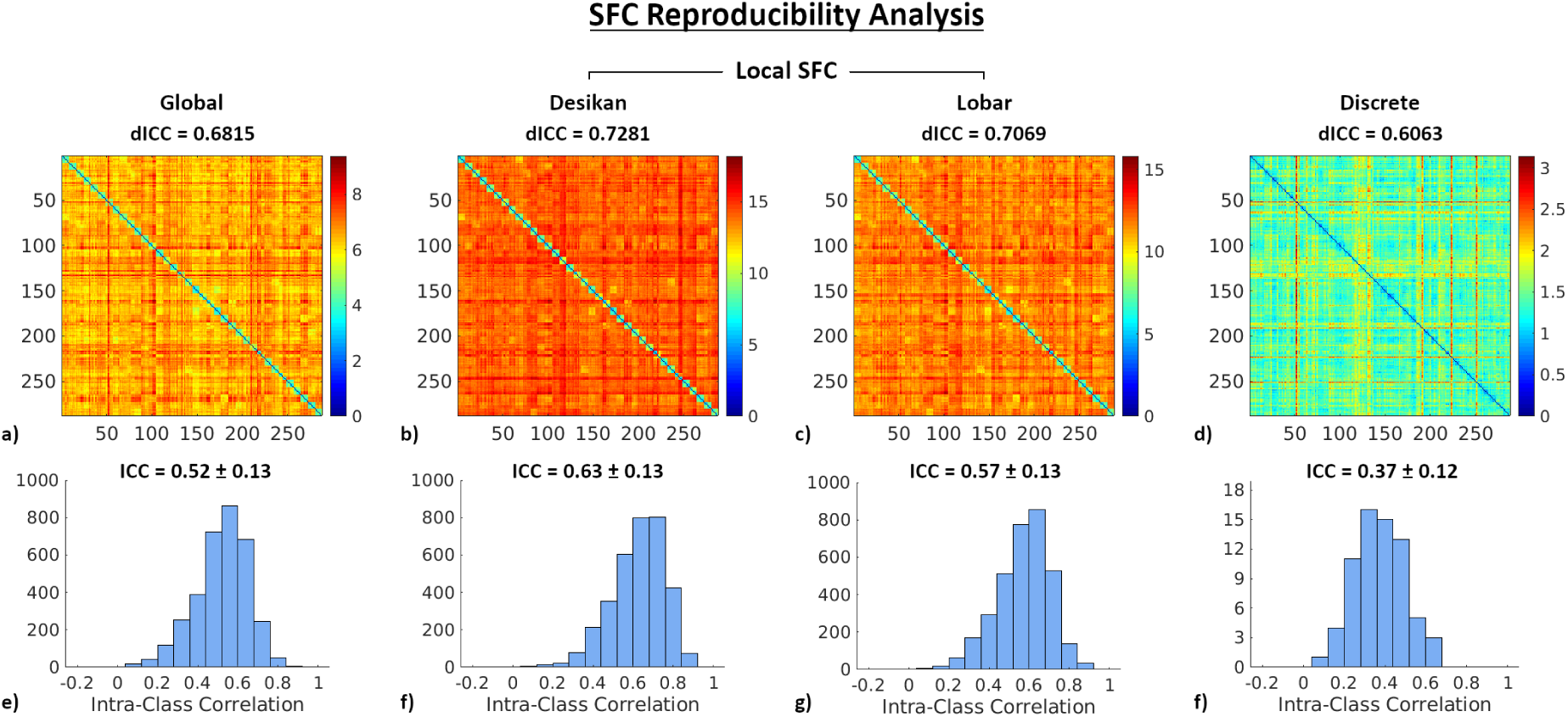
SFC reproducibility analysis. The first row from left to right shows pairwise distance matrices for *SFC*_*gbl*_, *SFC*_*locdk*_, *SFC*_*loclb*_, *SFC*_*dct*_, respectively. The second row shows the corresponding histograms of ICC at each node. dICC values are displayed above each distance matrix and the means and standard deviations are displayed above each histogram.

We also compare intra- and inter-subject *SFC*_*gbl*_ differences via paired t-tests. Figure 8 shows the −*log*(p-value) maps on the left hemisphere for four randomly selected pairs of subjects (first row) to show inter-subject comparisons and two randomly selected subjects (second row) to show intra-subject comparisons. Overall, we see more significant differences (larger negative log p-values) of *SFC*_*gbl*_ between subjects than within subjects. Quantitatively, on average, 20-25% of nodes have p-values less than 0.05 for intra-subject comparisons. These differences may be due to a few sources: (1) subject-specific variations of the structural and functional connectomes over time (test and retest) and (3) variations introduced by some processing steps in the SBCI pipeline, e.g., registration, segmentation, surface reconstruction, fiber tracking, etc. We find that 55 − 65% of nodes have p-values less than 0.05 for inter-subject comparisons, representing much larger differences of the *SFC*_*gbl*_ feature between subjects than within subjects.

**Figure 8:**
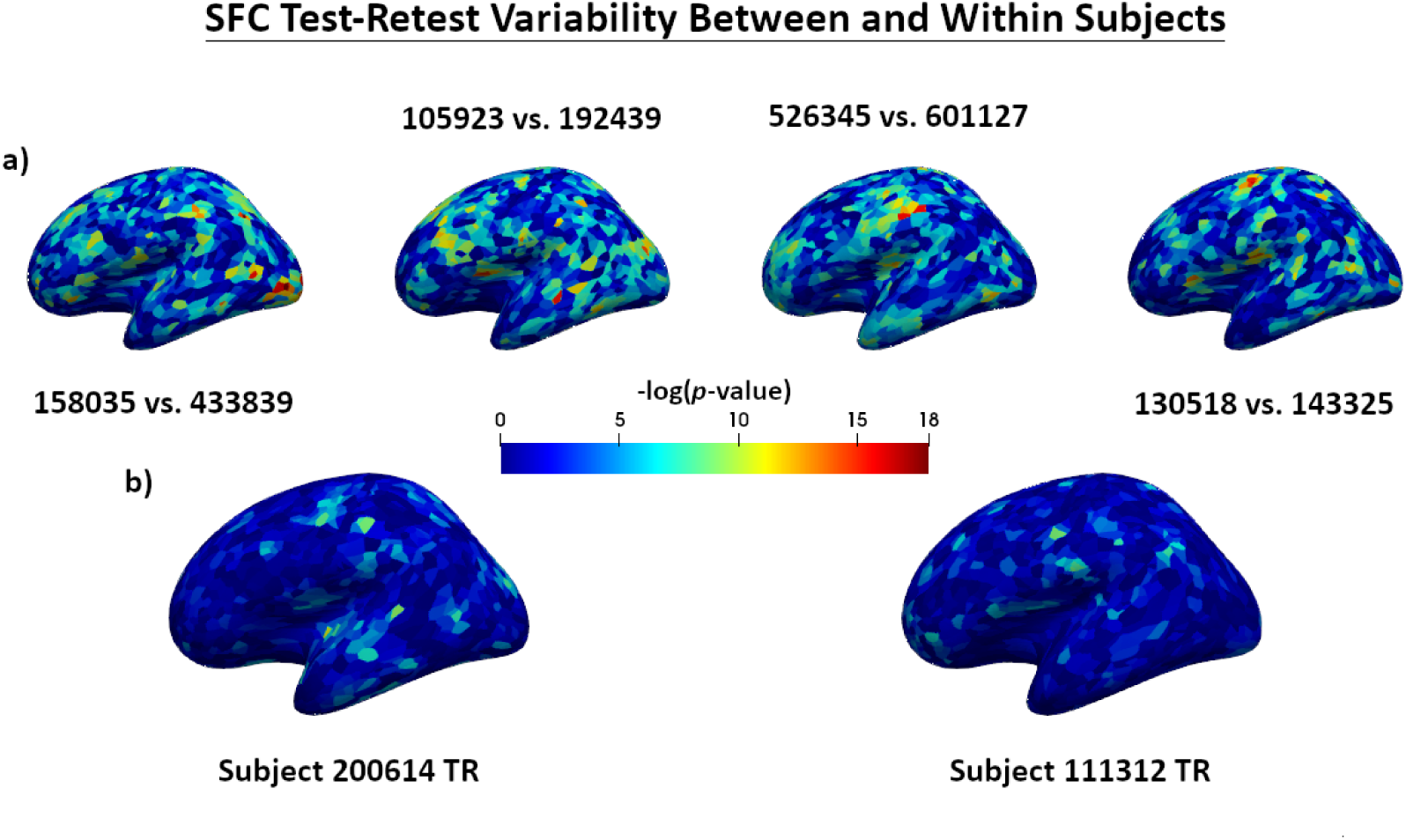
Negative log *p*-values of *t*-tests for the difference in *SFC*_*gbl*_ (a) between subjects, and (b) between the test-retest (TR) scans of individual subjects.

Our results show that the within subject *SFC*_*gbl*_, *SFC*_*locdk*_, *SFC*_*loclb*_ and *SFC*_*dct*_ are reproducible (although to different degrees) in data from two sessions that occurred in the span of several months in healthy young adults, and the *SFC*_*gbl*_ is more variable between subjects than within subjects. Therefore, we expect the SFC features to be informative and robust biomarkers to detect individual or group effects.

### 3.4. SC-FC Coupling Sex Difference

Finally, we evaluate the usefulness of our four SFC features (*SFC*_*gbl*_, *SFC*_*locdk*_, *SFC*_*loclb*_, and *SFC*_*dct*_) as biomarkers to detect group differences in sex. We use a small subset of the HCPYA data (due to high computational times and storage demand for each subject), containing 43 randomly selected males and 46 randomly selected females from the 26-30 year-old group in the S500 data release. Figure 9 shows the average SFC features across all males and females for the *SFC*_*gbl*_, *SFC*_*locdk*_, and *SFC*_*loclb*_ features on the inflated brain surface. Additionally, we display the −*log*(p-values) of the t-tests performed between the male and female cohorts in Figure 10.

**Figure 9:**
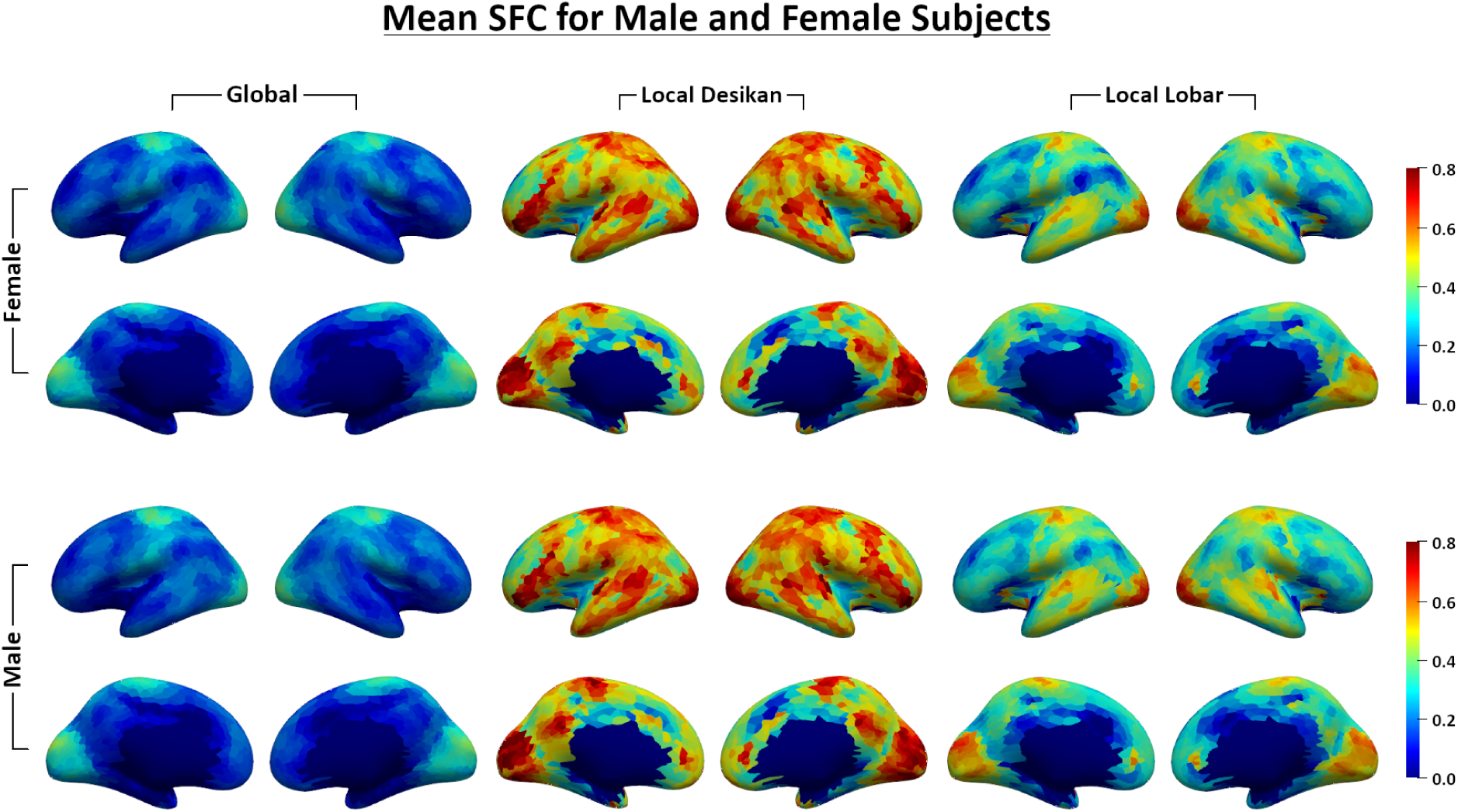
Mean *SFC*_*gbl*_, *SFC*_*locdk*_, and *SFC*_*loclb*_ values calculated over 43 male and 46 female subjects.

**Figure 10:**
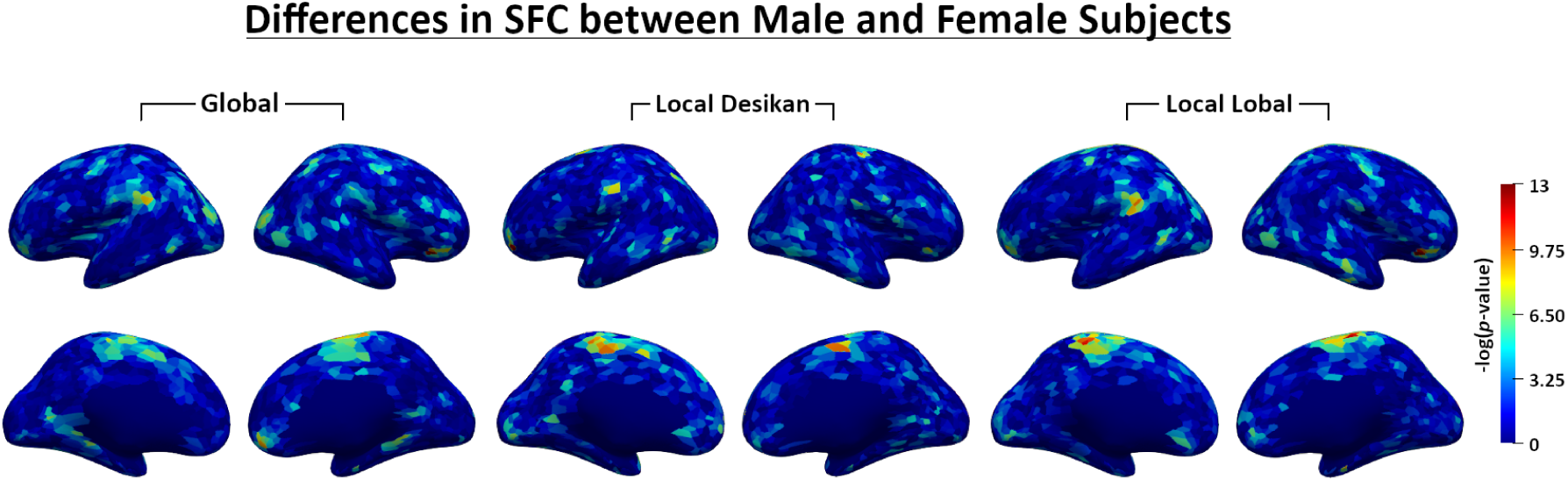
Negative log *p*-values of *t*-tests for the difference in SFCs between male and female subjects.

From the first two columns in Figure 9, we see that the global SC and FC are more correlated in primary sensory/motor areas such as S1/M1 and the visual and auditory cortices and less correlated in secondary association areas like the prefrontal cortex. The spatial distribution of *SFC*_*gbl*_ is consistent with the fundamental organizing architecture of the brain known since Brodmann’s map was published in 1909 [Brodmann, 2007]. Brodmann’s work began with the idea that “specific physiological functions in the cerebral cortex depend on specific histological structure and connectivity”. This principle is clear in primary sensory areas where specific histological patterns and cortical layer structures are closely associated with functional activity [Zeki, 2016]. More modern mapping identified the hierarchical map organization according to unimodal and multimodal association areas [Mesulam, 2000]. Our global SFC feature peaks in sensory areas and gradually drop into unimodal and multimodal association areas. These observations highlight the ability of our approach to advance brain mapping using modern data measurement techniques based on MRI. Moreover, comparing the *SFC*_*locdk*_ with the *SFC*_*gbl*_ shows that the *SFC*_*locdk*_ contains much higher values of structural-functional coupling, indicating that within ROI similarity is higher than whole brain similarity of SC and FC. The SFC decreases as we merge smaller ROIs in Desikan-Killiany into large lobar ROIs.

Figure 10 shows the −*log*(p-value) maps testing differences between males and females for the continuous SFCs on the surface. Sex differences were observed most strongly in the ventro-medial prefrontal cortex, the somatosensory-motor areas, the supra-marginal gyrus, and the occipitoparietal areas extending into the fusiform gyrus. These results emphasize how structure and function are differentially related between the sexes and are consistent with reported behavioral differences between men and women. For example, men and women perform differently on emotional recognition [Lausen and Schacht, 2018] and emotional decision making [van den Bos et al., 2013], which is well known to engage the ventro-medial prefrontal cortex [Bechara et al., 2000]. Men and women are also known to perform differently on facial recognition [Herlitz and Lovén, 2013], visual motion processing [Murray et al., 2018], and episodic memory recollection [Yonker et al., 2003]. Direct correlation between our measure of sex-related SFC differences and cognitive-behavioral differences between the sexes remains to be tested.

In addition to calculating group difference maps between males and females, we also try to predict sex from the continuous SC, continuous FC, and each of the four described SFC features. Table 2 shows the mean ± standard deviation of the ROC AUCs from each fold within a 5-fold stratified cross validation for each prediction model. PCA is performed first on each feature to reduce their dimensionalities to the top 5, 10, and 20 PCs. We then apply the logistic regression classifier (LRC), support vector classifier (SVC), and random forest classifier (RFC) to classify males and females.

Table 2 shows that the continuous SC is capable of predicting sex very well for 10 PCs, while the continuous FC is not as good at predicting sex. As for the SFC features, the continuous local Desikan-Killiany and local lobar SFCs have the best prediction scores, on par with the SC feature. The global SFC has higher predictive scores than FC, but not as high as SC nor the two continuous local SFC features. Interestingly, the discrete SFC from the Desikan-Killiany atlas has the worst prediction scores out of all of the models, even worse than many of the FC models.

## 4. Discussion

In this paper, we develop a surface-based connectivity integration (SBCI) framework and demonstrate that SBCI can extract reproducible and discriminative high-resolution structural (SC) and functional (FC) connectomes from high-quality MRI data on the white surface of the brain. Additionally, SBCI produces three subject-level imaging biomarkers that are reflective of the relationships between structural and functional brain signals. By using the HCP Test-Retest data, we show that SBCI can build reproducible SC and FC compared to previous connectivity studies [Honey et al., 2009; Noble et al., 2019] and three different structural-functional coupling features, two of which have not been described in previous literature to the best of our knowledge. Further, using data from the HCP Young Adult study, we demonstrate that these novel SFCs show good discriminative power as biomarkers. In summary, SBCI has the following unique advantages to other connectivity pipelines:

1. SBCI projects SC and FC to the white surface and does not rely on any brain parcellations. However, with a given parcellation, SBCI can easily transform the continuous SC and FC to discrete adjacency matrices.
2. SBCI represents both SC and FC in a continuous fashion on the white surface.
3. SBCI produces three novel imaging biomarkers reflecting the structural-functional relationships across the brain.

Most existing studies of SC and FC use GM volumetric ROIs, since volumetric ROIs provide convenient ways of grouping streamlines for SC and averaging BOLD signals for FC. However, GM ROIs can result in biased SC since streamlines can stop prematurely in the WM or near the GM-WM interface [Reveley et al., 2015; St-Onge et al., 2018]. Also, GM ROIs as nodes in FC often requires volumetric smoothing, reducing the spatial localization of BOLD signals [Coalson et al., 2018] and providing inaccurate FC estimation. Since structural signals are in the WM regions while functional signals are more present in the GM regions, we posit that the best place to study the integration of SC and FC is the white surface. Using the white surface to study FC is promoted by the HCP team [Coalson et al., 2018], however there are very few studies utilizing the white surface for SC study. The difficulties of studying SC on the white surface come from a few aspects: unreliable diffusive signals to infer WM fiber curves [Reveley et al., 2015], gyral biases due to low image resolution and tractography seeding [St-Onge et al., 2018], and huge computational burden in computing streamlines for constructing more reliable and high-resolution SC on the surface. SBCI uses a novel tractography algorithm [St-Onge et al., 2018] together with the KDE smoothing technique to overcome these challenges. More importantly, projecting both SC and FC to the white surface allows us to extend the traditional definitions of SC and FC to a continuous framework.

By treating SC and FC in a continuous fashion, we obtain high-resolution SC and FC by overcoming some of the typical limitations that arise from imaging acquisition and image processing, e.g., relying on a discrete brain parcellation to get brain connectivity matrices [Honey et al., 2009; Bastiani et al., 2012; Fornito et al., 2013; Finn et al., 2015; Zhang et al., 2018]. Others have previously treated SC in a similar manner [Moyer et al., 2017, 2016] and showed that continuous SC is reproducible. In this paper, we smooth the SC to obtain continuous SC and using the HCPTR dataset show that smoothing increases reproducibility at high resolutions. Further, by extending the typically sparse SC to be a continuous feature, we close the sparsity gap between SC and FC, allowing us to interrogate structural-functional relationships more robustly, as FC is naturally more dense. This continuous treatment of SC also allows us to overcome some of the computational challenges that come with big data; we do not need to recover hundreds of millions of streamlines in order to obtain reproducible SC at high resolutions (compare Supplementary Material Table 3 and Table 1).

Using predefined parcellations to calculate discrete adjacency matrices to study structural-functional coupling is a standard choice in the literature [Honey et al., 2009; Chamberland et al., 2017; Honey et al., 2010; Buckner et al., 2013; Ghumman et al., 2016; Baum et al., 2020; Cocchi et al., 2014; Jiang et al., 2019], with almost no exceptions. However, from our results in Figure 7 and Table 2, we see that the discrete SFC (derived from Desikan-Killiany atlas-based SC and FC) has several drawbacks: (1) it has low resolution (the number of ROIs in the Desikan-Killiany atlas); (2) it is less reproducible compared with our novel continuous local and global SFC features; and (3) it is less powerful in distinguishing groups. The continuous SFC features on the other hand overcome these drawbacks. But compared with the discrete SFC, one limitation of the continuous SFC features is that we need to perform registration for group-wise analyses. The registration in this paper is done using features derived from the T1 image after surface reconstruction from Freesurfer. A more straightforward registration based on the SC, FC, and SFC features is needed.

Three different SFC features, *SFC*_*gbl*_, *SFC*_*loc*_, and *SFC*_*dct*_, are proposed in this paper. The *SFC*_*gbl*_ characterizes how the SC between a given location and all other locations in the brain relates to the FC at that location and all other locations. In discrete space, this is analogous to the correlation between a row of the FC matrix and the corresponding row of the SC matrix. That is, both short range and long range connections to the vertex will contribute the final value of *SFC*_*gbl*_. On the other hand, *SFC*_*loc*_ measures the similarity of SC and FC within a predefined region, bounded as defined by a given atlas. From the results in Figure 9, we can see that at most vertices, *SFC*_*loc*_ is larger than *SFC*_*gbl*_, indicating that local SC and FC have much greater similarity within ROIs than global SC and FC. To test the discriminative ability of these biomarkers, we conduct a predictive analysis to classify sex. We use PCA to reduce the dimensionality of each biomarker before fitting various predictive models. Using 10 PC scores and the SVC model, we obtain high AUCs for all of the continuous SFC features. Interestingly, the local SFCs perform better than the global SFC in this particular task. However, note that we only used the Desikan-Killiany atlas to calculate *SFC*_*loc*_, which may not be an optimal atlas for studying SFC [Messé, 2019]. Experiments on different atlases should be conducted in the future.

One known limitation of the SBCI pipeline is that SET is sensitive to the input surfaces, according to our results from the HCPTR dataset. SET uses the geometry of the surface a priori to initiate WM streamlines. If surface reconstructions are significantly different across two scanning sessions for a single subject, then SBCI is unlikely to produce reliable SC for that subject. As such, we removed two outliers as determined by the differences in white surface reconstructions between the test and retest scans for our reproducibility analyses. Future work for improving SBCI should also focus on more robust surface reconstruction e.g., better segmentation and surface reconstruction methods [Zhao et al., 2019; Henschel et al., 2020], or collecting and incorporating high resolution T2w or FLAIR images to the T1w processing pipeline [Van Essen et al., 2013; Glasser et al., 2013; Renvall et al., 2016; Zaretskaya et al., 2018].

We have demonsrated that the SBCI pipeline can reliably reconstruct high-resolution SC, FC, and three novel SFC features. Future work should examine using SBCI to explore pathological differences in various neurological diseases, including HIV-associated cerebral small vessel disease (CSVD), mild cognitive impairment (MCI), Alzheimer’s disease (AD), and chronic pain, among others. Finally, we intend to use SBCI to determine an optimal brain parcellation for studying structural, functional, and structural-functional coupling for individual and group effects.

## 5. Acknowledgements

This research is partially supported by grant MH118927, MH118020 and AG066970 from the United States National Institute of Health.

Data were provided in part by the Human Connectome Project, WU-Minn Consortium (Principal Investigators: David Van Essen and Kamil Ugurbil; 1U54MH091657) funded by the 16 NIH Institutes and Centers that support the NIH Blueprint for Neuroscience Research; and by the McDonnell Center for Systems Neuroscience at Washington University. We also thank M.D. Paul Geha and Dr. Feng Vankee Lin for helping interpret these novel SFC features and their differences in sex.

