## Supplemental Material for "Surface-Based Connectivity Integration"

### 1. Local Data

As part of an ongoing study approved by the Research Subjects Review Board (RSRB) at the University of Rochester, both HIV-infected (HIV+) subjects and HIV-uninfected (HIV-) controls were recruited to study the effects of CSVD. Consent was obtained prior to any evaluation in accordance with RSRB approval. Twenty controls without signs of CSVD were selected for the development of the SBCI pipeline.

All imaging was conducted on a research dedicated 3T Siemens MAGNETOM Prisma (Erlangen, Germany) scanner equipped with a 64-channel head coil. High-resolution T1-weighted (T1w) anatomical images were acquired using the MPRAGE (magnetization prepared 3D rapid gradient echo) sequence with GRAPPA acceleration factor of 2 at the resolution of or up-sampled to 1 $mm^3$. Diffusion imaging was performed using a 2D transverse single-shot single-echo spin echo (SE) echo-planar imaging (EPI) sequence with 64 diffusion directions at two non-zero b-values (1,000 and 2,000 $s/mm^2$) each and 7 reference scans at 1.5 mm isotropic resolution. Parallel imaging was enabled using a slice acceleration factor of 3 and phase acceleration of 2 for a total scan time of 11 min 38 s. Functional imaging was performed using a 2D transverse single-shot gradient echo (GE) EPI sequence with TR=993 ms and echo spacing=4.43 ms at 2 mm isotropic resolution. Parallel imaging was enabled using a slice acceleration factor of 8 for a total scan time of 5 min 9 s. Reverse phase encoding gradient field maps were also acquired for distortion correction.

### 2. Full Image Preprocessing Steps

Diffusion image preprocessing steps include brain extraction using BET [Smith, 2002], susceptibility induced distortion correction using TOPUP [Andersson et al., 2003], and eddy-current induced distortion and subject motion corrections using EDDY [Andersson and Sotiropoulos, 2016]. The eddy corrected images are then postprocessed using Mrtrix3 [https://www.mrtrix.org/], ANTs [Avants et al., 2011], and the Sherbrooke Connectivity Imaging Lab toolbox in python (Scilpy). Next, the diffusion images are skull stripped, bias-field corrected, cropped, intensity normalized, and resampled to 1 mm isotropic resolution. The $b = 0$ images are extracted for coregistration with the T1w image. To prepare for tractography, we calculate the fiber orientation distribution function (fODF).

The anatomical T1w image is registered to the high-resolution average $b = 0$ diffusion image using ANTs before being processed by Freesurfer's recon-all pipeline [http://freesurfer.net/], which includes motion correction, intensity normalization, skull stripping, neck removal, tissue segmentation,

[1]Equal contribution.

*601 Elmwood Avenue, Rochester, NY, USA, 14642

surface mapping and inflation, spherical mapping, cortical parcellation to the Desikan and Destrieux atlases, and curvature and cortical thickness calculations and statistical analyses.

Functional MRI data preprocessing is carried out using the Freesurfer Functional Analysis Stream (FsFast) tools. Briefly, the following procedures are applied: motion correction using the first volume as the refrence image (mc-sess), smoothing using a 3D Gaussian kernel with FWHM 5 mm (spatialsmoothing-sess), slice-timing correction using Fourier-space time-series phase-shifting (stc-sess), non-brain removal and brain-mask creation (mkbrainmask-sess), registration to the T1w in high-resolution diffusion space using boundary-based registration (BBR) [Greve and Fischl, 2009], resampling to the surface (rawfunc2surf-sess) and surface smoothing with FWHM 5 mm (surfsmooth-sess). Surface resampling is performed onto the left and right hemispheres of the anatomical image in the native diffusion space.

### 3. Parameter Selection in SBCI

#### 3.1. Seed Count

| Atlas | Steps | $N_s = 1.5$M | $N_s = 3$M | $N_s = 4.5$M | $N_s = 6$M | $N_s = 7.5$M |
|---|---|---|---|---|---|---|
| Desikan | $t = 75$ | 0.7460 | 0.7469 | 0.7470 | 0.7471 | 0.7470 |
| | $t = 100$ | 0.7467 | 0.7474 | 0.7473 | 0.7475 | 0.7476 |
| Destrieux | $t = 75$ | 0.7505 | 0.7513 | 0.7515 | 0.7515 | 0.7516 |
| | $t = 100$ | 0.7498 | 0.7511 | 0.7514 | 0.7515 | 0.7515 |
| Atlas-Free | $t = 75$ | 0.7245 | 0.7306 | 0.7328 | 0.7338 | 0.7345 |
| | $t = 100$ | 0.7241 | 0.7296 | 0.7314 | 0.7323 | 0.7328 |

Table 1: dICC vs. Seed count (in millions (M) per hemisphere). The table shows results for three approaches using two values for the SET parameter $t$ (surface flow size).

Table 1 shows the results of SET (without smoothing) varying the number of seeds and value of the flow parameter $t$ on the HCPTR data. The dICC values are calculated using two atlases (Desikan-Killiany and Destrieux) and the Atlas-Free approach. Table 2 shows the same analysis after removing two subjects determined to be outliers due to large differences in white surface reconstructions between the test and retest scans.

| Atlas | Steps | $N_s = 1.5$M | $N_s = 3$M | $N_s = 4.5$M | $N_s = 6$M | $N_s = 7.5$M |
|---|---|---|---|---|---|---|
| Desikan | $t = 75$ | 0.7789 | 0.7797 | 0.7799 | 0.7801 | 0.7800 |
| | $t = 100$ | 0.7803 | 0.7811 | 0.7811 | 0.7812 | 0.7813 |
| Destrieux | $t = 75$ | 0.7794 | 0.7803 | 0.7805 | 0.7806 | 0.7806 |
| | $t = 100$ | 0.7783 | 0.7796 | 0.7799 | 0.7800 | 0.7801 |
| Atlas-Free | $t = 75$ | 0.7376 | 0.7443 | 0.7467 | 0.7478 | 0.7485 |
| | $t = 100$ | 0.7372 | 0.7430 | 0.7450 | 0.7460 | 0.7466 |

Table 2: dICC vs. Seed count (in millions (M) per hemisphere) after removing outliers. The table shows results for three approaches using two values for the SET parameter $t$ (number of steps to compute the flow).

Due to the amount of time, computational power and storage needed to run the tractography algorithm, we only explore two values of the flow parameter $t$. From prior experience, we select $t = 75$ and $t = 100$ as reasonable values for the analysis. The flow parameter $t = 75$ produces higher dICC values than $t = 100$, and thus more reproducible SC, for both the Destriuex and Atlas-Free approaches regardless of the number of seeds used. Additionally, for all approaches, dICC increases as a function of the number of seeds used, but this increase is not appreciable enough to justify the increased computational power and storage needed when using more seeds. Removing the outliers increases the dICC values for all three approaches, particularly for the Desikan-Killiany and Destrieux approaches.

### 4. SC-FC Coupling

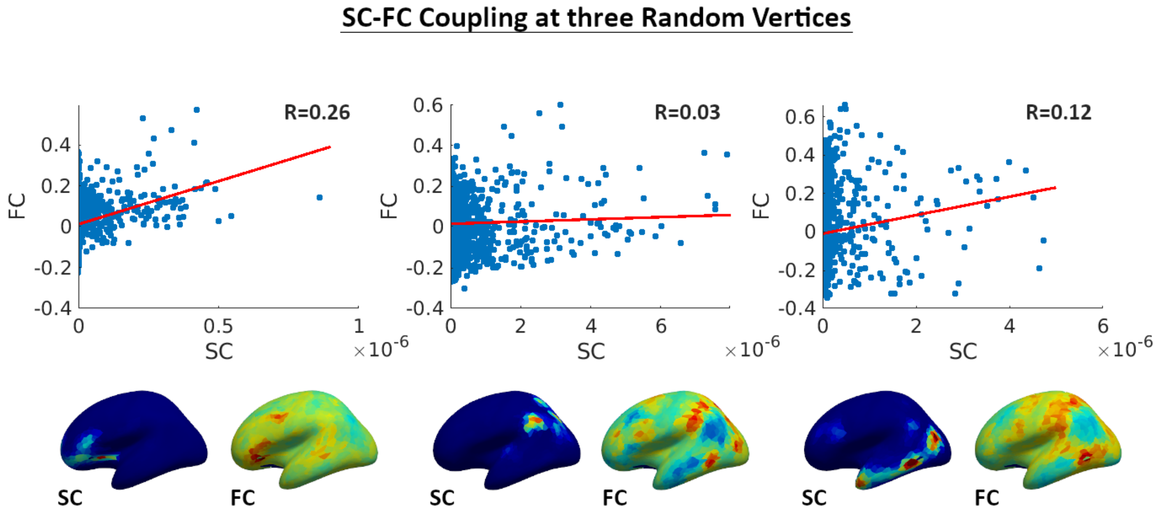


Figure 1: Seed based connectivity of SC and FC on a random subject using a single random vertex as the seed for each plot. The scatter plots show the correlation between SC and FC for each of the seeds with the corresponding connectivity plotted on the inflated surface underneath.

### 5. Connectivity Resolution

Figures 2 and 3 show the connectivity matrices and global SFC for varying resolutions for one random subject from the HCPTR data and one random subject from the CSVD data. The FC matrices produced from the lower imaging resolution of the CSVD data look comparable to those produced using the HCPTR data, displaying similar block patterns. Note, however, that there appears to be more noise in the CSVD FC matrices. This noise likely comes from only having 5 min of resting state data in the CSVD study, compared to the 15 min runs collected for HCP. The SC matrices obtained from both data sets also look comparable. The grid obtained after removing 95% of the vertices from the surface and the grid after removing 99% of the vertices appear to produce very similar overall patterns of the connectivity matrices.

Similarly, figures 4 and 5 show that the global SFC calculated using grids obtained from removing 95% and 99% of the vertices on the full surface mesh is comparable at the two resolutions. Additionally, there appear to be similar patterns in SFC between the two data sets, e.g. high values of SFC in the occipital lobes.

### 6. Discrete SFC

Figure 7 shows the discrete SFC calculated on the Desikan-Killiany atlas projected to the inflated surface. Interestingly, the discrete approach appears to produce slightly different patterns of SFC when compared to global and local SFC, e.g. lower SFC in the occipital lobe, higher SFC in some areas of the frontal lobe.

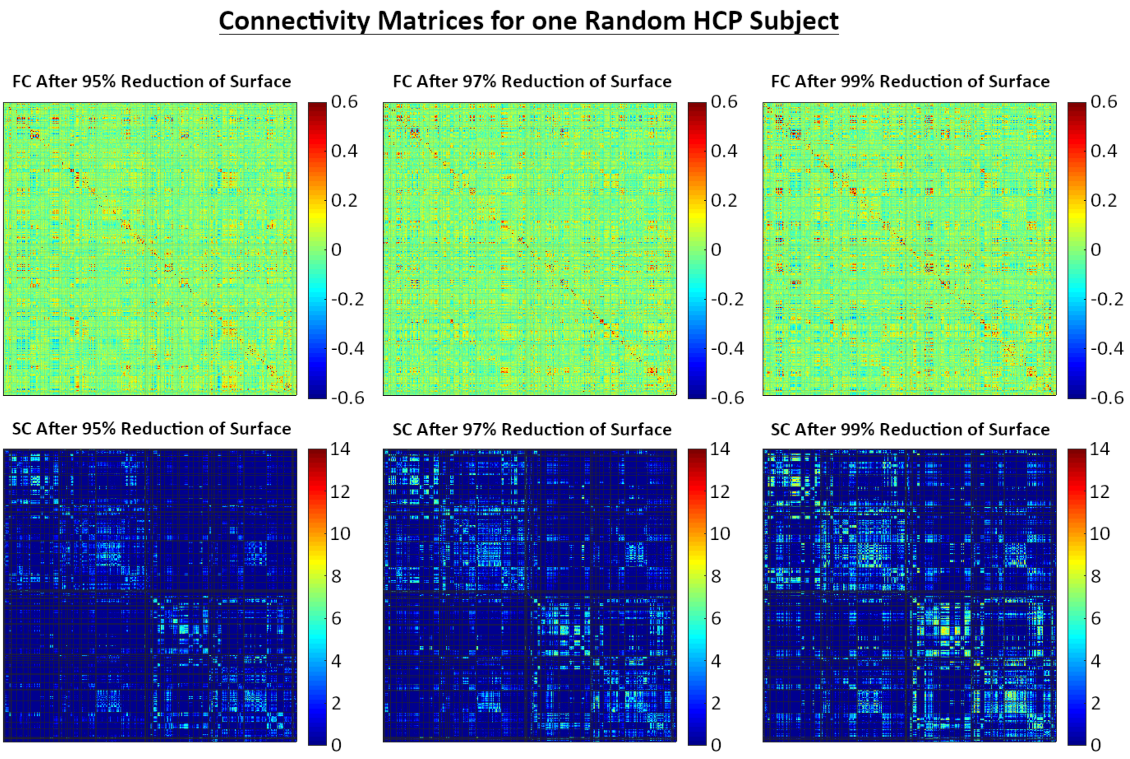


Figure 2: Connectivity matrices for one random HCP subject at three different resolutions.

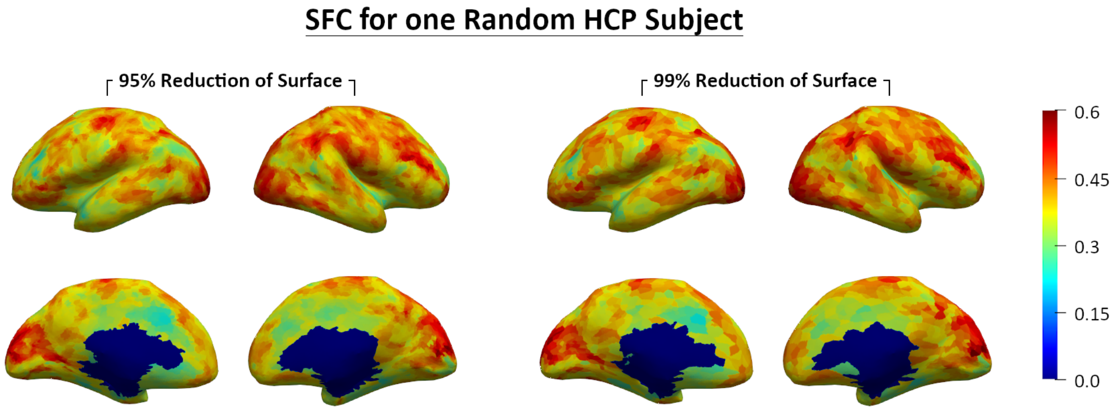


Figure 3: The global SFC values for one random HCP subject plotted on the inflated surface at two resolutions. SFC values are calculated on grids of around 12,000 and 4000 vertices obtained by reducing the total number of vertices on the surface by 95% and 99%, respectively.

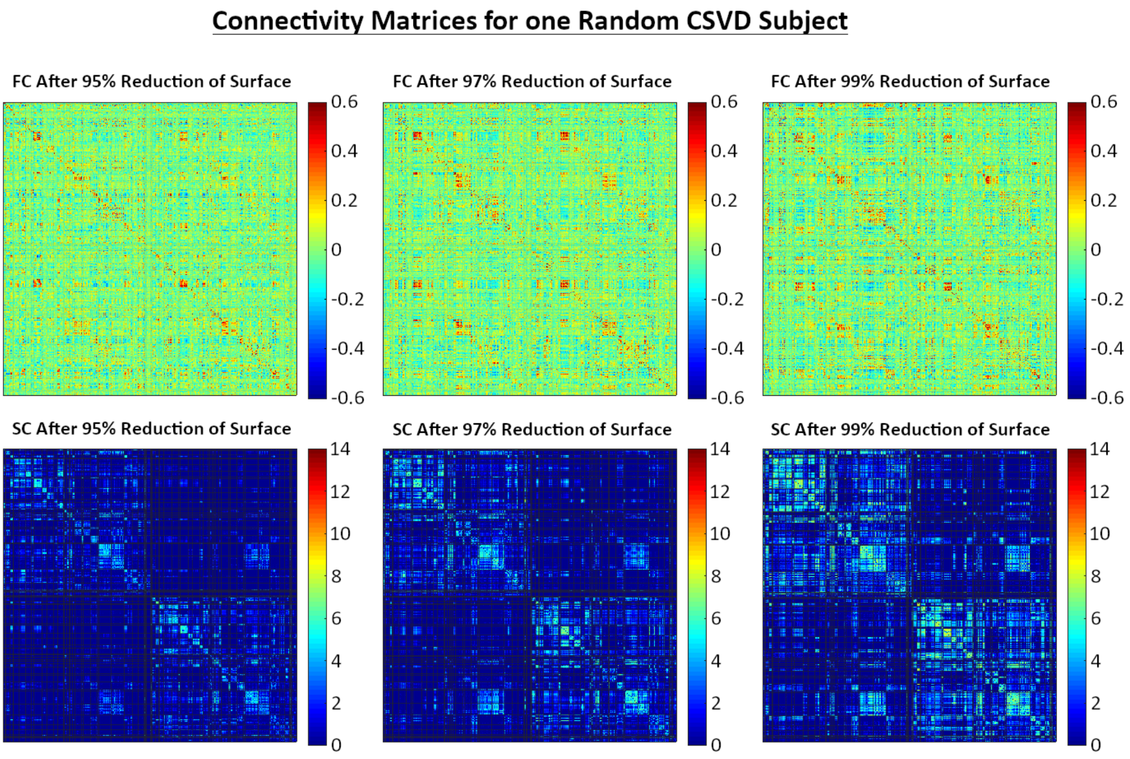


Figure 4: Connectivity matrices for one random CSVD subject at three different resolutions.

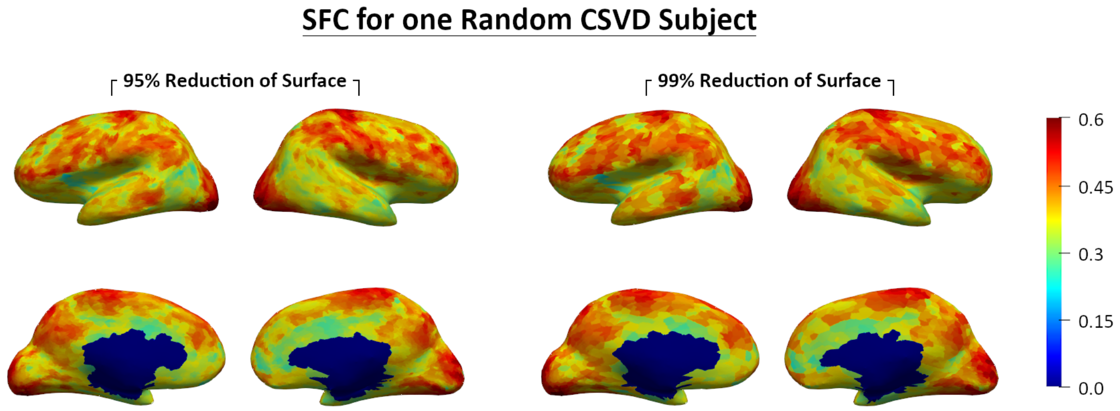


Figure 5: The global SFC values for one random CSVD subject plotted on the inflated surface. SFC values are calculated on grids of around 14,000 and 5000 vertices obtained by reducing the total number of vertices on the surface by 95% and 99%, respectively.

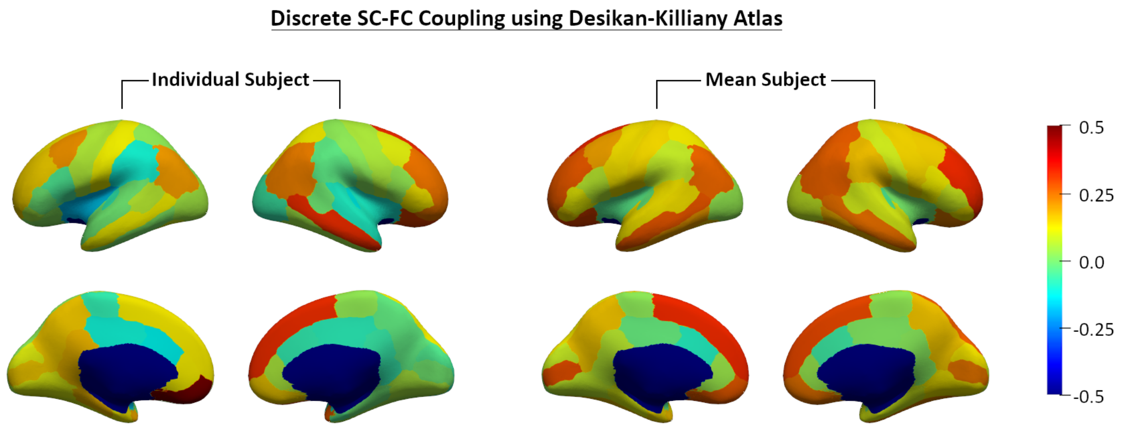


Figure 6: The discrete SFC values for one random HCPTR subject and the mean SFC values from all HCPTR subjects that we used plotted on the inflated surface.

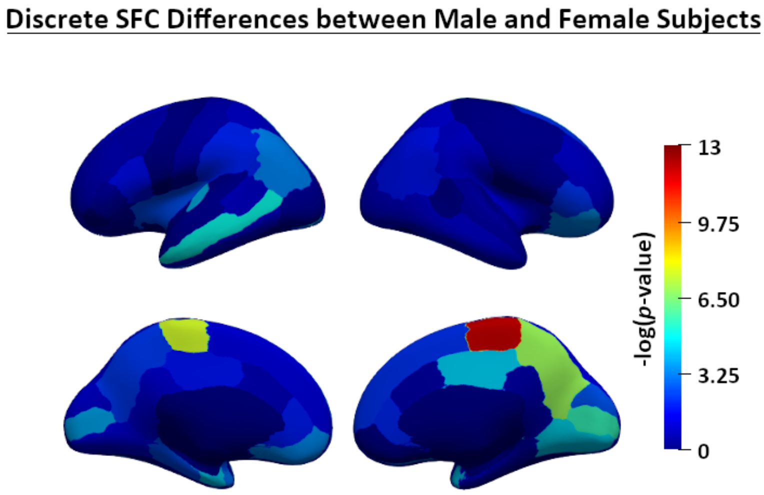


Figure 7: Negative log $p$-values of $t$-tests for the difference in discrete SFCs between male and female subjects using the Desikan-Killiany Atlas.
